# Anchoring of actin to the plasma membrane enables tension production in the fission yeast cytokinetic ring

**DOI:** 10.1101/586792

**Authors:** Shuyuan Wang, Ben O’Shaughnessy

## Abstract

The cytokinetic ring generates tensile force that drives cell division, but how tension emerges from the relatively disordered ring organization remains unclear. Long ago a muscle-like sliding filament mechanism was proposed, but evidence for sarcomeric order is lacking. Here we present quantitative evidence that in fission yeast ring tension originates from barbed-end anchoring of actin filaments to the plasma membrane, providing resistance to myosin forces which enables filaments to develop tension. The role of anchoring was highlighted by experiments on isolated fission yeast rings, where sections of ring unanchored from the membrane and shortened ~30-fold faster than normal [Mishra M., et al. (2013) *Nat Cell Biol* 15(7):853-859]. The dramatically elevated constriction rates are unexplained. Here we present a molecularly explicit simulation of constricting partially anchored rings as studied in these experiments. Simulations accurately reproduced the experimental constriction rates, and showed that following anchor release a segment becomes tensionless and shortens via a novel non-contractile reeling-in mechanism at about the load-free myosin-II velocity. The ends are reeled in by barbed-end-anchored actin filaments in adjacent segments. Other actin anchoring schemes failed to constrict rings. Our results quantitatively support a specific organization and anchoring scheme that generates tension in the cytokinetic ring.

---

Cytokinesis is the final stage of the cell cycle, when cells assemble an actomyosin contractile ring that constricts the cell into two (Pollard and O’Shaughnessy, 2019). The basic mechanical function of the ring is to generate tension, and hence to exert inward force that drives cell cleavage. Ring tensions of 10 – 50 nN were reported in echinoderm embryos (Rappaport, 1967; Yoneda and Dan, 1972; Hiramoto, 1975; Rappaport, 1977) and ~ 400 pN in protoplasts of the fission yeast *Schizosaccharomyces pombe* (Stachowiak *et al*., 2014), but the mechanism that produces tension is not established.

How do the molecular components coordinate to generate tension in the ring? Following its discovery, and the finding that actin and myosin-II are major components, the parallels with muscle were compelling and a muscle-like sliding filament mechanism was proposed (Schroeder, 1972; Fujiwara and Pollard, 1976; Mabuchi and Okuno, 1977). However, contractile rings appear relatively disordered and evidence is lacking for the highly ordered architecture of muscle based on the sarcomere repeat unit (Green *et al*., 2012; Laplante *et al*., 2015). A second, conceptually simple possibility is that ring tension derives from passive elasticity, using similar principles to those governing elastic fibers in the extracellular matrix (Kielty *et al*., 2002).

A third possible mechanism, consistent with the disorder seen in contractile rings, is based on the anchoring of individual actin filaments to the plasma membrane by their barbed ends. No sarcomeric organization or permanent crosslinking between actin filaments is involved. Rather, overlapping filaments randomly positioned around the ring are independently made tense by myosin-II that binds and pulls each filament. The net ring tension is the sum of such independent filament contributions. This mechanism of tension production in individual filaments was demonstrated during assembly of the ring of the fission yeast *S. pombe*, when anchored myosin-II pulls barbed-end-anchored actin filaments and renders them tense so as to draw together the node precursors of the ring (Vavylonis *et al*., 2008). Super-resolution studies suggest that anchoring of actin filament barbed ends to the plasma membrane persists in constricting fission yeast rings, suggesting this tension mechanism persists (Laplante *et al*., 2016), and further support is provided by molecularly detailed simulations of constricting fission yeast rings that reproduce experimentally measured ring tensions (Stachowiak *et al*., 2014; O’Shaughnessy and Thiyagarajan, 2018; Pollard and O’Shaughnessy, 2019).

This last mechanism requires firm lateral anchoring of barbed ends to the membrane, to oppose the myosin force and allow the filament to become tense. Thus, if anchoring were compromised the ring tension would be severely impacted. The effect of anchor loss was examined by Mishra et al. (Mishra *et al*., 2013) in an experimental study of fission yeast cell ghosts, permeabilized cells that lack cytoplasm and provide a unique laboratory to study isolated cytokinetic rings. On addition of ATP, entire sections of rings became unanchored by pulling away from the plasma membrane (Fig. 1). Subsequently, the unanchored segments shortened until they became taut, whereas anchored segments did not visibly contract. The shortening rate of the unanchored segments, 0.22 ± 0.09 μm s^−1^, was ~30-fold the normal constriction rate and was independent of initial ring length.

**Fig. 1.**
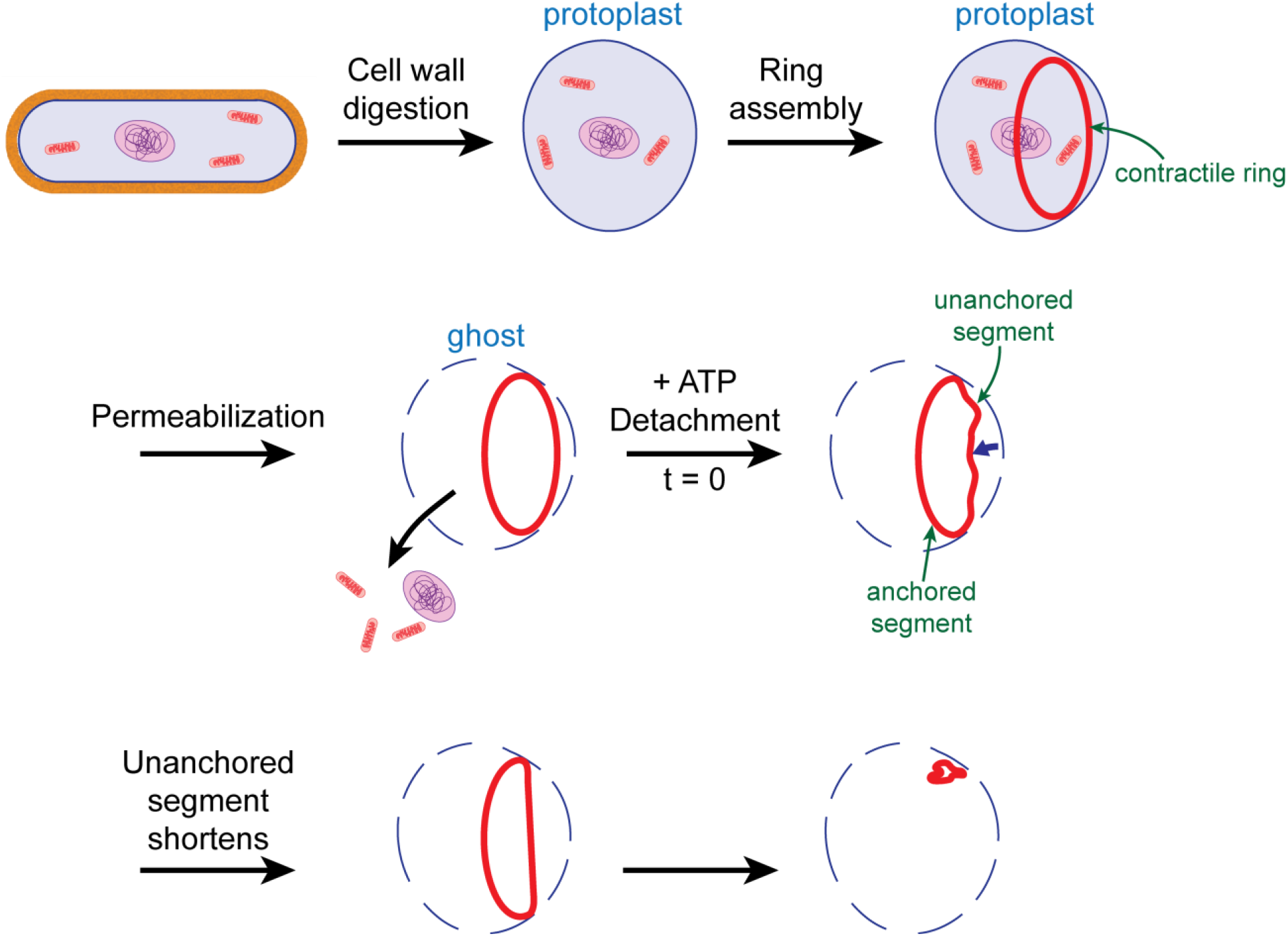
Schematic of cytokinetic ring constriction observed in permeabilized fission yeast protoplasts by Mishra et al. (Mishra *et al*., 2013). Protoplasts were generated from normal intact yeast cells by digestion of the cell wall, a fraction of which assembled cytokinetic rings. The plasma membrane was then permeabilized by detergent treatment so that cytoplasm escaped, resulting in cell ghosts that contained isolated contractile rings, lacking the highly viscous cytoplasm and its constituents. On addition of ATP, in a typical sequence a segment of ring detached from the weakened membrane and then shortened at ~30-fold the normal constriction rate until it became straight. While the unanchored segment shortened, the anchored segment remained of fixed length. Subsequent detachment and shortening sequences complete ring constriction. Note that during the shortening and straightening episode the unanchored segment is dragged through the aqueous medium, whose viscosity is presumably far less than that of the cytoplasm.

The findings of Mishra et al. (Mishra *et al*., 2013) are unexplained, and the implications for the ring tension mechanism are not established. Here, we mathematically model constriction of partially anchored cytokinetic rings, comparing the predictions to a series of measurements reported in ref. (Mishra *et al*., 2013). We begin with a simple argument to show that these experimental findings are inconsistent with either a passive elastic or a sarcomeric muscle-like tension mechanism. We then develop a molecularly detailed mathematical model of the fission yeast ring which shows that the results of ref. (Mishra *et al*., 2013) are in quantitative agreement with the barbed-end-anchoring tension hypothesis. The model shows that tensionless unanchored segments shorten by a novel non-contractile “reeling-in” mechanism, and accurately reproduces the observed shortening rate which is identified as the load-free velocity of myosin-II.

## Results

### Experimental shortening rates of unanchored ring segments are inconsistent with elastic or sarcomeric tension mechanisms

In the experiments of Mishra et al., segments of contractile rings in fission yeast ghosts became unanchored from the plasma membrane. The unanchored segments had a typical initial radius of curvature *R* ~2 μm and were pulled with velocity *ν* ~ 200 nm s^−1^ through the cell ghost aqueous contents as they shortened (Fig. 1). As the cytoplasm is removed in ghosts, the expected viscosity of the aqueous medium is similar to that of water, *ν*_water_ = 0.001 Pa·s. Thus the drag coefficient per unit length of a typical ring segment of length *L* ~2 μm and thickness *w*~ 0.2μm (Stachowiak *et al*., 2014) is approximately (Broersma, 1960) *ζ~* 4π*ν*_water_ /[ln(2*L/w*) – 0.31] ~10^−3^ pN μm^−1^. Balancing viscous drag and tensile forces, *T/R* ~ ζν, yields a negligible tension *T* ~2 × 10^−3^ pN, some 5 orders of magnitude smaller than the ~ 400 pN reported experimentally (Stachowiak *et al*., 2014).

Thus, unanchored segments have essentially zero tension. This conclusion is inconsistent with a passive elastic mechanism, since an elastic ring segment would retain its tension even when unanchored. Further, the shortening rate was independent of the initial length of the unanchored segment (Mishra *et al*., 2013), inconsistent with a sarcomeric mechanism, for which the rate would be proportional to the number of unanchored sarcomeres *n*_sarc_ and hence the initial segment length. Indeed, the experimental shortening rate is of order the load-free myosin-II velocity, 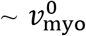, whereas we find that an unanchored sarcomeric ring segment would shorten much more rapidly, at 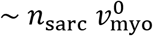 (see below).

### Model of the fission yeast cytokinetic ring and its application to partially anchored rings

If unanchored ring segments have zero tension, why do they shorten? And why is the shortening rate ~ 0.22 μm s^−1^, ~30-fold the normal rate? To address these issues quantitatively we developed a molecularly detailed 3D simulation of the *S. pombe* cytokinetic ring anchored to the inside of the plasma membrane, Fig. 2. The *S. pombe* ring is particularly well characterized, as the amounts and biochemical properties of many contractile ring proteins were measured (Wu and Pollard, 2005; Arasada and Pollard, 2014; Goss *et al*., 2014), severely constraining the model (see Supplementary Note 1 and Table S1 for model details and parameters). The formin-capped actin filaments (Kovar *et al*., 2006) are anchored at their barbed ends to the membrane (Laplante *et al*., 2016). Anchored myosin-II clusters (Schroeder and Otto, 1988; Naqvi *et al*., 1999) exert force according to the force-velocity relation for myosin-II (Edman, 1979), and bind to and pull upon actin filaments dynamically crosslinked by α-actinin dimers (Fig. 2a). Lateral mobilities of anchors in the membrane were previously determined from component velocities measured in live cells (Stachowiak *et al*., 2014).

**Fig. 2.**
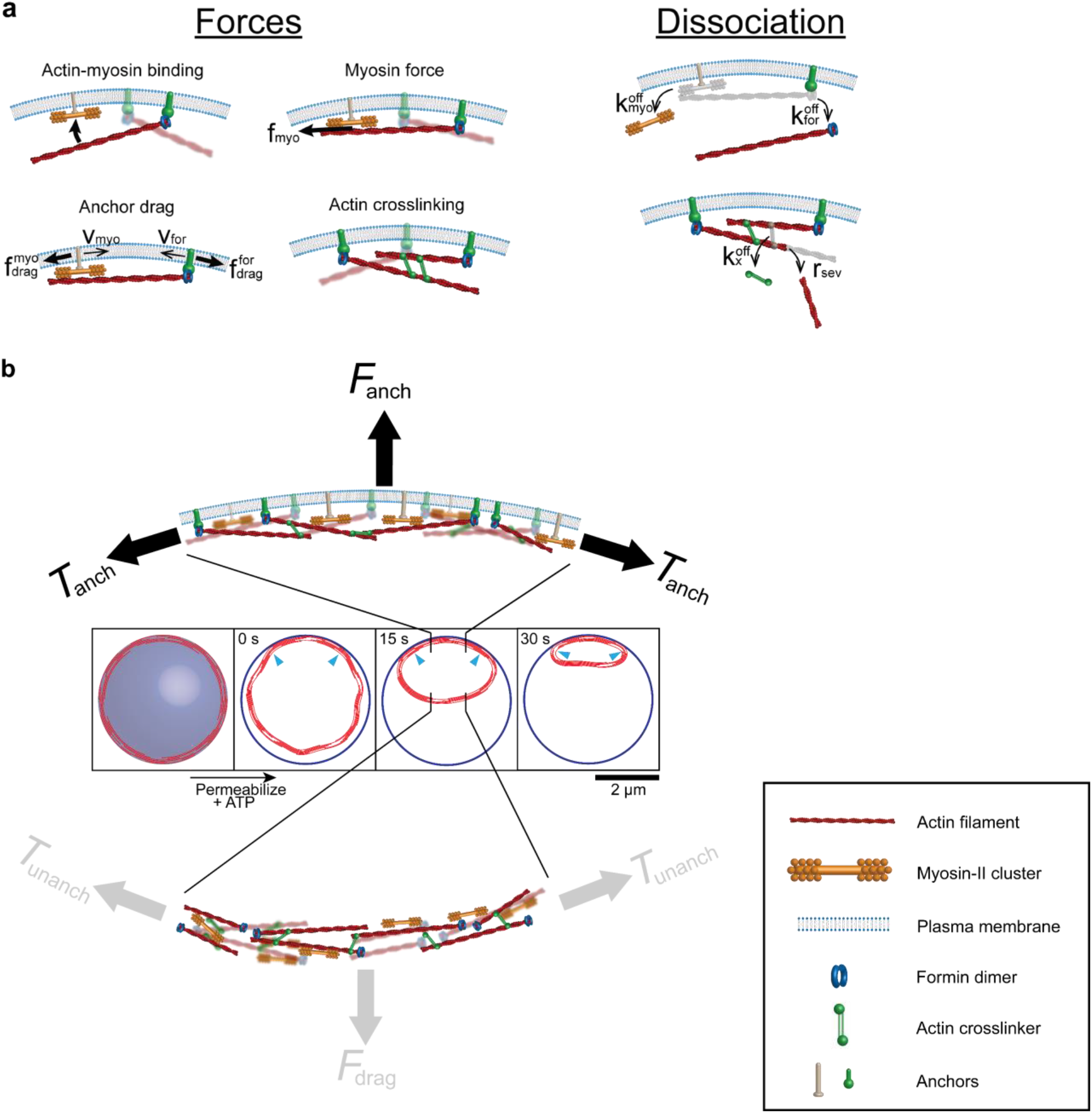
Barbed end anchored 3D model of the cytokinetic ring in permeabilized fission yeast protoplasts (components not to scale). For component amounts and parameters see Supplementary Note 1 and Table S1. **(a)** Actin filaments, membrane-anchored by formin Cdc12p, bind membrane-anchored myosin-II that pulls actin following a linear force-velocity relation. Anchors move laterally, resisted by membrane drag forces, while drag from the aqueous medium acts on moving actin, myosin-II and formin. These components dissociate without replenishment, being absent from permeabilized cells (Mishra *et al*., 2013). Simulations were run without α-actinin crosslinkers, as they dissociated within ~ 2s; simulated constriction rates were unaffected (Fig. S2). Actin dissociates by unbinding with formins, and by stochastic cofilin-mediated filament severing. **(b)** Constriction of a partially unanchored ring. Initial ring lengths, 12-19 μm (Mishra *et al*., 2013). The initially anchored ring is a disordered bundle (**Stachowiak *et al*., 2014**). At t = 0 s (ATP addition), partial detachment is simulated by removal of anchors in a segment and a small displacement. Depicted ring shapes are from a simulation with an initial ring 17 μm long and 80% unanchored. The unanchored segment shortens, not the anchored segment. Arrowheads: anchored/unanchored interfaces. Top: a portion of the anchored segment (schematic). The ring tension *T*_anch_ balances the net force from anchors that attach components to the membrane, *F*_anch_. Bottom: In the unanchored segment, components can move in any direction. The ring tension is negligible, *T*_unanch_ ≪ *T*_anch_, because it balances a tiny net drag force from the aqueous medium, *F*_drag_.

The model is fully 3D and dynamic. The ring can follow any contour in space, and detailed positions and configurations of components are represented over the ring cross section and along its length (Fig. 2). Actin filaments, for example, can orient in any direction and assume any 3D shape, determined by the forces exerted upon them and the known bending stiffness of F-actin. Crowding effects are accounted for by interactions between components. Component motions are tracked in all directions; e.g., when a ring segment detaches from the membrane the components experience forces tending to pull them away from the membrane through the aqueous medium, while viscous drag forces from the medium oppose this motion (Fig. 1, 2b). The model can describe fast component motions and high constriction rates, essential to capture the 30-fold normal constriction rates in cell ghosts. The model autonomously constricts the ring, as the ring length is continuously updated according to the evolving component locations.

We used the model to simulate constriction of partially unanchored rings in permeabilized protoplasts (Figs. 1, 2b). We begin with a normal steady state ring, a ~ 0.2 μm wide bundle of randomly positioned myosin-II clusters and actin filaments oriented parallel to the ring with random polarity, consistent with experiment and simulations of fully anchored rings (Stachowiak *et al*., 2014) (Supplementary Note 1). At time *t* = 0, the myosin-II and formin anchors are removed in a segment of the ring, mimicking an initial detachment episode following ATP addition (Mishra *et al*., 2013). As most cytoplasmic constituents are absent in cell ghosts (Mishra *et al*., 2013), binding of new components (Pelham and Chang, 2002; Yonetani *et al*., 2008; Stachowiak *et al*., 2014) is absent. Dissociation rates were determined from the experimentally measured loss in component amounts in ghosts over the course of constriction and are considerably reduced from normal (Mishra *et al*., 2013) (Table S1).

Component velocities in the unanchored segment are computed by balancing aqueous medium viscous drag forces with myosin-II and other forces. A similar force balance describes the anchored segment, but components are confined to the membrane and experience membrane anchor drag forces. The ring tension is computed by summing actin filament tensions over the cross section.

### Unanchored ring segments in simulations are tensionless

Using initial ring lengths of 12-19 μm (Mishra *et al*., 2013), we compared simulated ring shapes, constriction rates and tensions to experiment (Mishra *et al*., 2013). In a typical simulated tension profile 1 s after detachment (Fig. 3a) the tension is negligible in the unanchored segment, but substantial in the anchored segment.

**Fig. 3.**
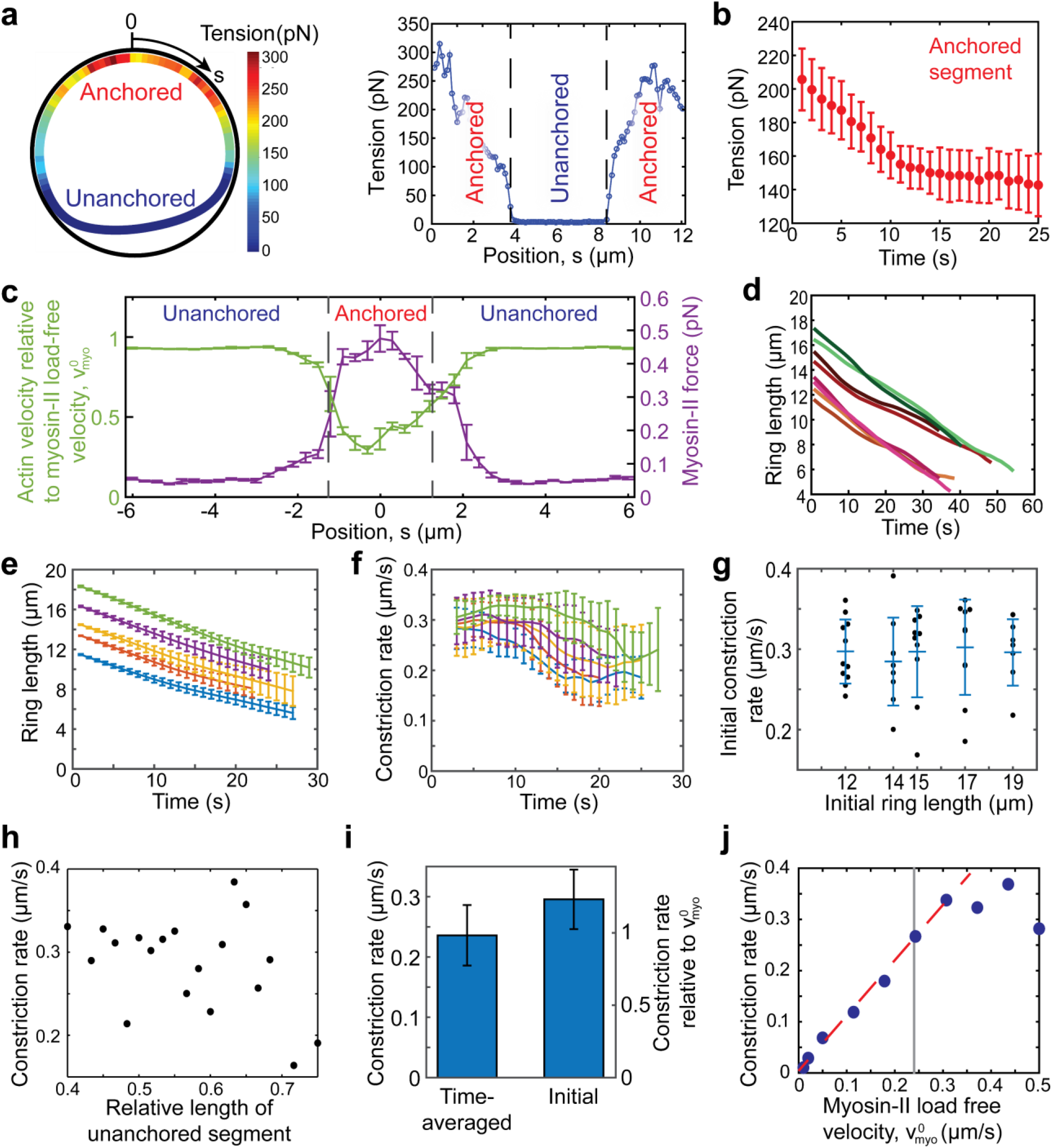
Unanchored ring segments have zero tension and constrict at close to the load-free velocity of mysoin-II, 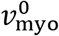, independently of initial length. Simulations of partially unanchored rings. Initial length 12.6 *μ*m, unanchored segment 20% of ring length, unless otherwise stated. Other parameters, see Supplementary Note 1 and Table S1. Error bars: s.d. **(a)** Anchoring is required for ring tension. Tension in a simulated ring 1s after detachment of a segment 40% of ring length. The unanchored segment has almost zero tension. **(b)** Tension versus time in the anchored segment (n=10 simulations). 40% of ring initially unanchored. **(c)** Actin filament velocities relative to the myosin-II clusters they interact with (green) and mean forces exerted by myosin-II clusters per actin filament they interact with (purple). Mean values over a length 0.3 μm, one simulation. **(d)** Length of partially unanchored ring versus time for 8 simulations, initial lengths 12-18 μm. **(e)** Length of partially unanchored ring versus time for initial lengths 12, 14, 15, 17 and 19 μm averaged over n = 11, 9, 10, 10 and 7 simulations, respectively. **(f)** Mean constriction rates (rates of decrease of ring length) versus time for constrictions of **(e)**. **(g)** The initial constriction rate is independent of initial ring length (p = 0.96, one-way ANOVA). Constriction rates from **(e)** at 1s. Bars: mean ± s.d.. **(h)** The time averaged constriction rate and initial length of unanchored segment relative to total ring length are uncorrelated (correlation coefficient *r* = −0.40, *p* = 0.094, n=19 rings). **(i)** Time averaged and initial constriction rates averaged over all constrictions of **(e)**, n = 47. **(j)** In simulations with a range of 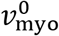 values, the time-averaged constriction rate was 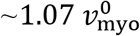 over most of the range (95% confidence interval: 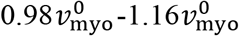, least-squares fit of first 7 points, dashed line). Gray line: value used throughout this study (0.24 *μ*m s^−1^).

In the anchored segment the tension peaked at a mean value 342 ± 51 pN (n = 10), similar to the experimentally reported ~ 390 pN for normally anchored rings (Stachowiak *et al*., 2014). With time, tension in the anchored segment decreased due to component dissociation and incoming actin filaments that saturated anchored myosin-II clusters (Fig. 3b).

Thus, the model reproduces the almost vanishing tension of unanchored ring segments in experiments. The origin of the anchoring requirement for tension is apparent from the ~ 0.5 pN force that myosin-II clusters exerted on barbed end-anchored actin filaments they interacted with in the anchored region (Fig. 3c), sustained by large anchor drag forces (10.7 ± 5.0 pN at 10 s, *n* = 321 filaments in 10 simulated rings). This created tension in the slowly moving filaments. By contrast, the unanchored segment was tensionless because myosin slid unanchored actin filaments at close to the load-free velocity, 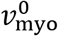, working against almost zero drag force (Fig. 3c).

### The model reproduces experimental constriction rates that are independent of ring length

In simulations, in agreement with experiment, the unanchored segments shortened but not the anchored segments. The simulated constriction rates were remarkably close to the experimental values. Despite being tensionless, simulated unanchored segments shortened until they became almost straight after ~ 25-60 s (Fig. 2b, 3d,e and Movie S1). The shortening rate was independent of the initial length of ring or unanchored segment, and approximately constant in time, with a mean time average 0.24 ± 0.05 μm s^−1^ (n = 47), Figs. 3e-i. Anchored segments scarcely shortened (~ 3% shortening, Movie S2). These results reproduce the observations of ref. (Mishra *et al*., 2013), where only unanchored segments shortened and the constriction rate, 0.22 ± 0.09 μm s^−1^, was consistent over many cells with rings of variable initial length.

### Unanchored ring segments are non-contractile and are reeled in at their ends

Remarkably, the shortening of unanchored ring segments was not contractile, revealed by the constant separation between fiducial markers in simulated constricting rings (Fig. 4a). One might imagine the rapid shortening of tensionless unanchored segments is driven by a contraction mechanism working against zero load, similar to zero load muscle contraction (Edman, 1979). On the contrary, these segments shortened by being *reeled in* at their two ends where they joined the anchored segment. Each end was reeled in at about half the myosin-II load-free velocity, 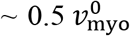, the mean velocity with which myosin entered the anchored segment (Fig. 4b,e and Movie S3), giving a net shortening rate 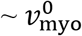. The non-contractile shortening left the myosin density in the unanchored segment constant in time and approximately spatially uniform (Figs. 4a,c), while reeled-in myosin accumulated in puncta of growing amplitude near the anchored/unanchored interfaces (Fig. 4a, arrowheads, and Fig. 4c).

**Fig. 4.**
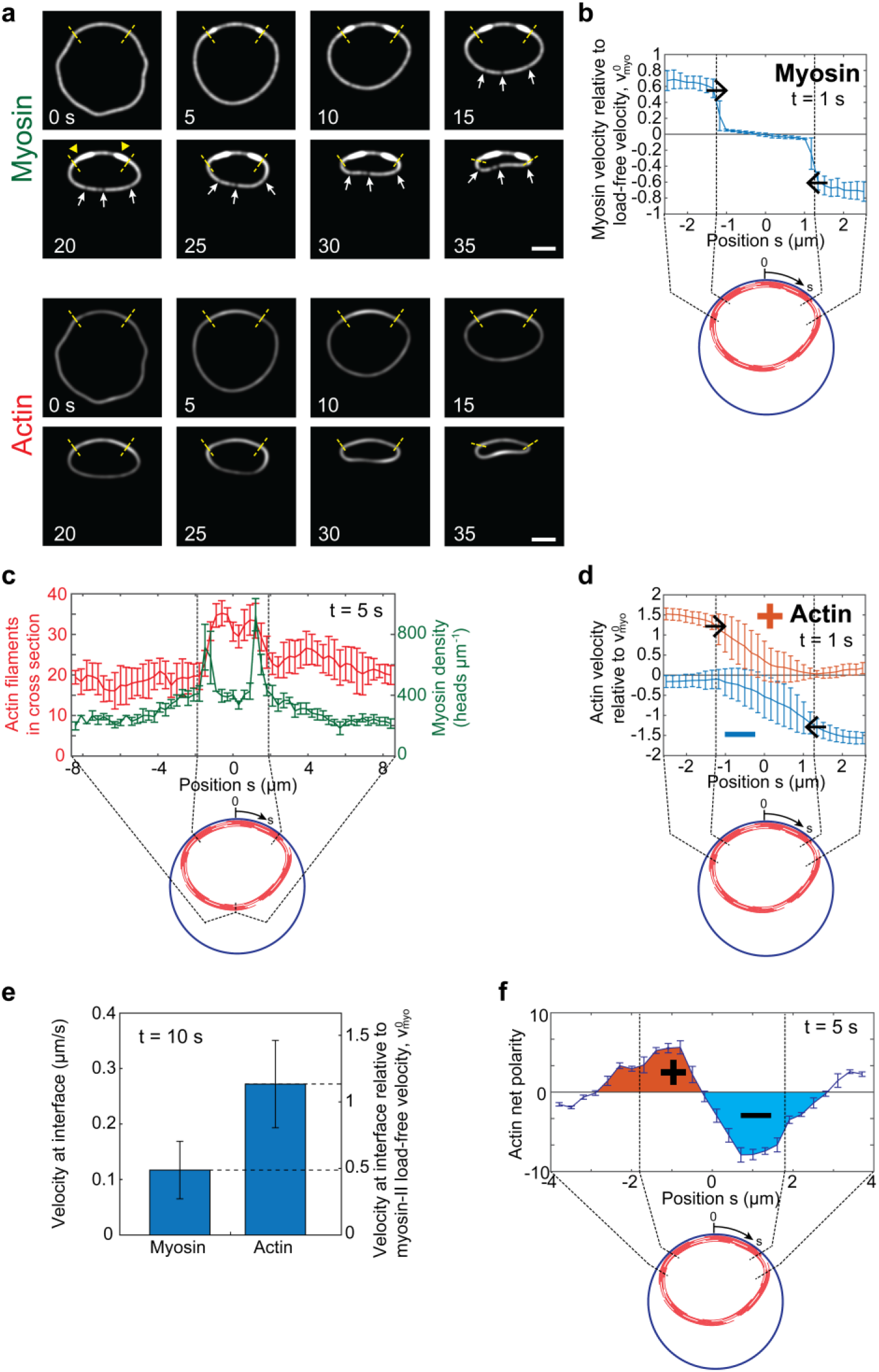
Unanchored ring segments are non-contractile and shorten by being reeled in at close to the myosin-II load-free velocity, 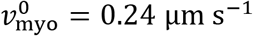. Simulation parameters, as for Fig. 3 unless otherwise stated. Error bars: s.d.. **(a)** Simulated time-lapse images of the constricting ring of Fig. 2b, with fluorescently tagged myosin-II and actin. Shortening of the unanchored segment is non-contractile: fiducial markers have constant separation (arrows), and myosin and actin densities remain constant. Instead, the unanchored segment is reeled in so that myosin-II accumulates in puncta (arrowheads) near the anchored/unanchored interfaces (dashed lines). The anchored segment does not shorten. Fluorescence imaging simulated with a 2D Gaussian point spread function, width 200 nm, centered on myosin-II clusters or actin subunits. **(b)** Velocity profiles of myosin-II in the interfacial and anchored zones after 1s. Unanchored myosin-II enters the anchored segment with velocity 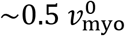, the reeling in velocity. Mean velocities parallel to the ring, averaged over a bin size 1/6 μm and 10 simulations. **(c)** Myosin and actin density profiles 5 s after partial detachment. Myosin puncta develop near each interface and actin accumulates in the anchored segment. Both densities are approximately uniform in the unanchored segment. Mean densities, averaged over a bin size 0.3 μm and n = 7 rings, initial length 19 μm. **(d)** Velocity profile of actin subunits belonging to clockwise-oriented (+) and anticlockwise-oriented (-) filaments in the interfacial and anchored zones after 1s. Unanchored filaments of a definite polarity enter the anchored region at each end, with velocity 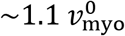. **(e)** Mean velocities of incoming unanchored myosin-II and actin as the components enter the anchored region, time 10s. The myosin velocity, 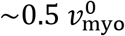, is the reeling-in velocity. The actin velocity, 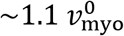, is less than 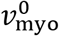 greater than the myosin velocity due to sliding resistance from anchored myosin clusters and interfacial crowding. Mean values over a region within 0.1 μm of the interface (60 myosin-II clusters, 543 actin subunits, 10 simulations). **(f)** Net actin polarity (number of clockwise-minus anticlockwise-oriented filaments) in the interfacial and anchored zones in simulations of **(c)**. Clockwise (anticlockwise) polarity bias develops near the left (right) interface.

### Rings constrict at close to the load-free velocity of myosin-II in permeabilized protoplasts

It is noteworthy that for both simulations and experiments the constriction rates are of order the myosin-II load-free velocity in our simulations, 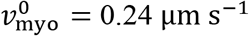 (Table S1). We stress that this is an experimental value, taken from ref. (Stark *et al*., 2010) where 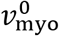 was measured in *S. pombe* rings versus number of myosin-II (Myo2p) molecules per actin filament which we set to 20 from the ratio of Myo2p to formin Cdc12p molecules measured in the ring in ref. (Wu and Pollard, 2005) (Supplementary Note 1). That the constriction rate could be related to 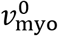 is plausible, since unanchored segments encounter negligible viscous drag force while shortening. To test this, we artificially varied 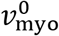 through the range 0.01-0.5 μm s^−1^. Simulations showed that the constriction rate 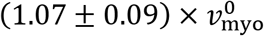 can indeed be identified as a constant times 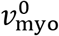 over a large range (Fig. 3j).

### Reeling in is caused by barbed-end anchored actin filaments

What causes reeling in, and why is the shortening rate close to 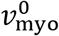? We found that reeling in is a direct consequence of the barbed end anchoring of actin filaments that is the basis of tension generation in normally anchored rings. The agents of reeling in are actin filaments in the interfacial zone whose barbed ends are anchored to the membrane in the anchored segment (Fig. 5c). About half of these filaments straddled the interface, their pointed ends oriented into the unanchored segment (Supplementary Note 1 and Fig. S1). These filaments grabbed unanchored myosin-II clusters and reeled in the unanchored segment at the load-free myosin velocity 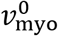, as the segment offered negligible load due to the low medium viscosity in cell ghosts. This process occurs at both ends, suggesting a net shortening rate 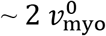. In practice, the rate 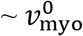 is somewhat lower (Fig. 3i, j), due to sliding resistance from anchored myosin clusters on incoming actin filaments and myosin crowding at the interface (Fig. 4 and Supplementary Note 1).

**Fig. 5.**
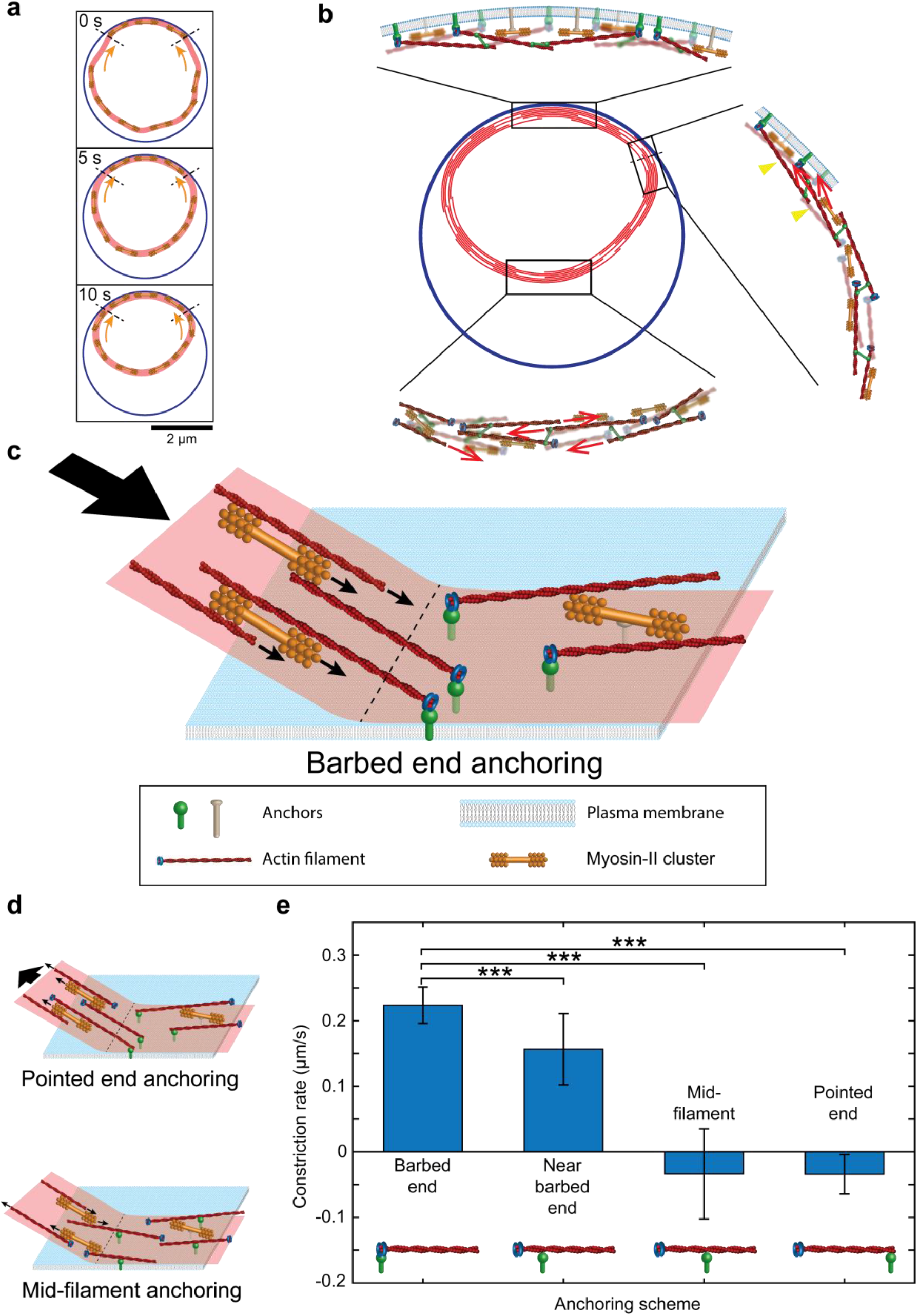
Reeling in mechanism of ring constriction in permeabilized protoplasts. **(a)** Unanchored ring segments are reeled in at their ends. The simulated ring of Fig. 2b is shown at the indicated times. Reeling in (arrows) at the interface with the anchored segment (dashed lines) is not contractile, so that on average the distance between myosin-II clusters remains constant (myosin-II clusters shown schematically, not to scale). **(b)** The three distinct zones of simulated partially anchored rings. In the unanchored segment (bottom), almost stationary myosin clusters propel actin filaments at the zero load velocity 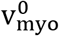 (arrows) clockwise or anti-clockwise depending on filament polarity. Components have much lower velocities in the anchored segment (top), due to viscous drag forces from component anchors in the plasma membrane. In the interfacial region (right) actin filaments and myosin clusters are reeled into the anchored segment at velocities of order 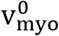 (arrows) by anchored actin filaments that bridge the interface and orient into the unanchored segment (arrowheads). **(c)** The reeling-in mechanism relies on barbed end anchoring of actin filaments to the plasma membrane. Filaments lie parallel to the ring, randomly oriented clockwise or anticlockwise. Thus, about half of those filaments anchored close to the anchored/unanchored interface (dashed line) straddle the interface, with their pointed ends oriented into the unanchored segment. These filaments grab unanchored myosin clusters and reel in the unanchored ring segment against almost zero load. This figure was adapted from (Pollard and O’Shaughnessy, 2019). **(d)** Pointed end or mid-filament actin anchoring schemes do not constrict rings in permeabilized protoplasts. With pointed end anchoring, actin filaments straddling the interface have the wrong orientation for reeling in, since myosin-II migrates to actin filament barbed ends. Instead, the unanchored segment is pushed outwards. Mid-filament anchoring produces zero net reeling in, as filaments of both orientations straddle the interface. **(e)** Simulated constriction rates for different anchoring schemes. Only anchoring at or near barbed ends constricts rings, and only barbed-end anchoring reproduces the experimental constriction rate of 0.22 μm s^−1^. Mean constriction rates, time-averaged over 20s (or until ring fracture, for pointed end anchoring) and over n=10 simulations. Model parameters as for Fig. 3. Error bars: s.d.

### Other actin anchoring schemes cannot reproduce experiment

That the model reproduces the experimental length-independent shortening rate of 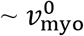 strongly supports the specific actin anchoring scheme assumed, at or near barbed ends. Other anchoring schemes cannot explain the experiments: with pointed end or mid-filament anchoring, actin filaments at the anchored/unanchored interface are wrongly oriented, and simulations failed to constrict unanchored segments (Fig. 5d, e). For example, with pointed end anchoring those anchored filaments that straddle the interface are oriented with barbed ends extending into the unanchored segment; since myosin-II migrates toward barbed ends, the unanchored ring segment tends to get pushed out rather than contract. With mid-filament anchoring, both orientations occur equally often (barbed or pointed ends, respectively, extending into the unanchored ring segment) so that contractile and expansive forces cancel.

### Constricting rings in permeabilized protoplasts have 3 distinct zone types

Thus, partially anchored rings have three distinct zone types (Figs. 5a,b). (1) In the anchored region anchored myosin clusters interacting with randomly oriented anchored actin filaments have small net velocity (Fig. 4b), giving a very small shortening rate. (2) The unanchored segment is a freestanding random actomyosin bundle. As drag forces are negligible, myosin works against zero load, generates no contractility or tension, and propels actin filaments with velocity 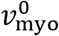 clockwise or anticlockwise (depending on polarity) that enter the anchored segment with a velocity 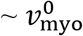 greater than the myosin reeling in velocity (Figs. 4d,e). These filament motions leave the actin density unaffected except for the latest stages (Figs. 4a,c), and produce net polarity puncta in the interfacial zones (Fig. 4f). (3) The interfacial zones, where non-contractile reeling-in occurs (Fig. 5a).

### Anchoring is required for ring constriction

The simulations showed that anchoring is required not only for tension, but also for ring constriction. Entirely unanchored simulated rings had almost vanishing tension and did not constrict (Fig. 6a, b and Movie S4), consistent with the images of constricting rings in ref. (Mishra *et al*., 2013) that featured at least one anchored segment. This is because reeling in of an unanchored segment requires the presence of an adjacent anchored segment whose barbed-end-anchored actin filaments execute the reeling in. Actin filament anchoring is the key requirement, as constriction occurred even without myosin anchoring in the anchored segment (Fig. 6a, b and Movies S5, S6).

**Fig. 6.**
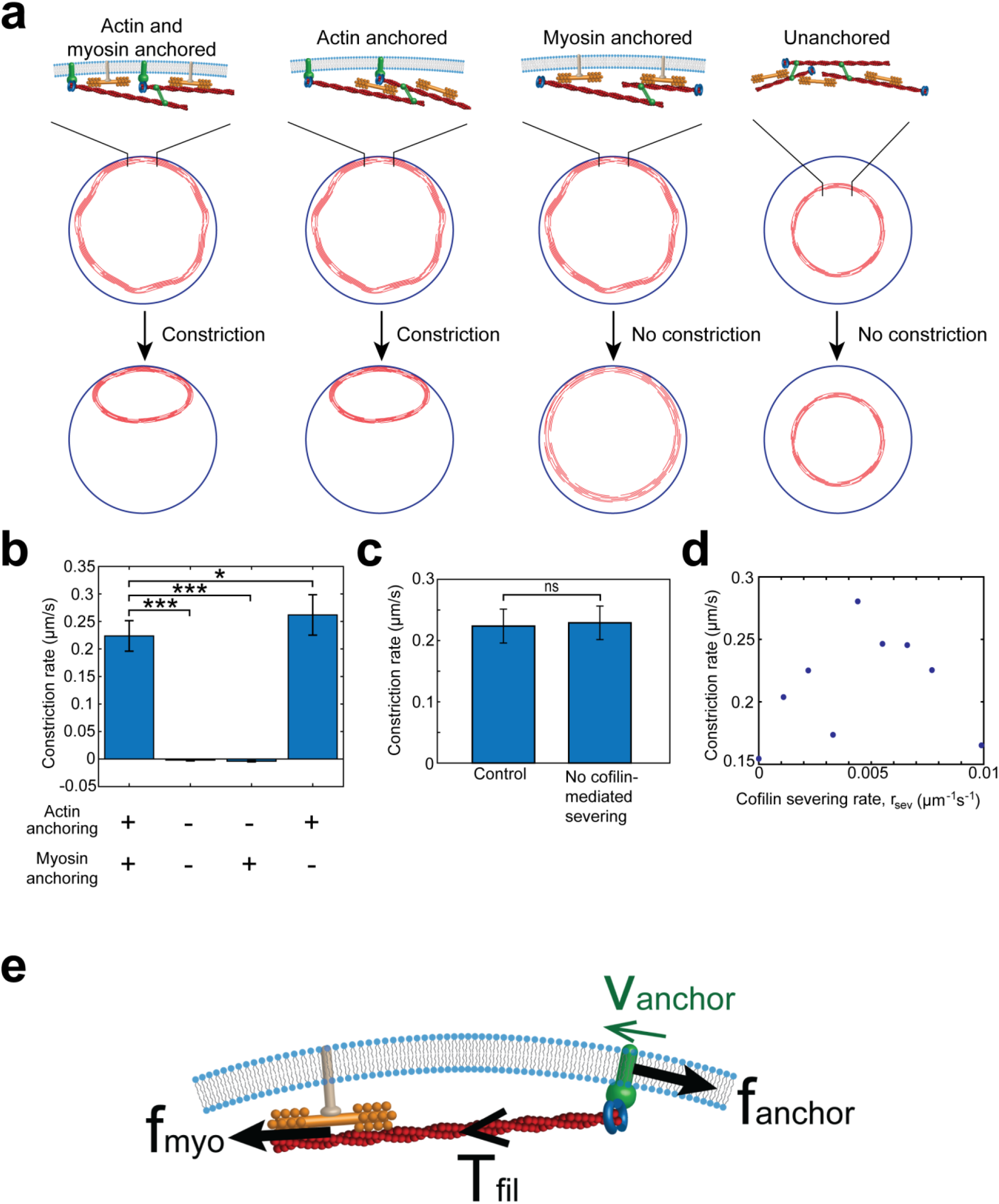
Constriction of cytokinetic rings in permeabilized protoplasts requires anchoring of actin but not actin turnover. Simulation results, model parameters as for Fig. 3. Constriction rates are from linear fits to simulated ring lengths versus time. **(a, b)** Partially anchored rings constrict when both actin and myosin are anchored (constriction rate 0.24 ± 0.05 μm s^−1^, Fig. 2c) or when only actin is anchored (constriction rate 0.26 ± 0.04 μm s^−1^). Loss of actin anchoring abolishes constriction, either for rings with only myosin anchored in the anchored segment (elongation rate 4±1 nm s^−1^) or for completely unanchored rings (elongation rate 2 ± 2 nm s^−1^). Mean values shown, averaged over n=10 (with actin anchoring) or n=13 (without actin anchoring) simulations. Error bars: s.d.. **(c)** There is no statistically significant difference between constriction rates of rings with (control) and without cofilin-mediated severing of actin filaments. Simulations without severing mimic the addition of jasplakinolide in the experiments of ref. (Mishra *et al*., 2013). Error bars: s.d.. **(d)** Constriction rate versus rate of cofilin-mediated severing of actin filaments. The constriction rate and severing rate are uncorrelated (n = 9, correlation coefficient *r* = 0.17, *p* = 0.65). **(e)** Schematic of anchoring mechanism for tension generation in the fission yeast contractile ring. A typical actin filament barbed end is anchored to the membrane via formin Cdc12p (blue) and a membrane anchor (green). The anchor moves laterally in the membrane when pulled by myosin-II, resisted by drag force *f*_anchor_. The myosin force and the resultant filament tension *T*_fil_ are substantial provided the anchor velocity *ν*_anchor_ is much less than the load-free myosin-II velocity 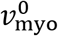.

### Dependence of fission yeast myosin-II activity on ATP concentration

The load-free velocity, 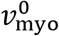, is a fundamental molecular property of myosin-II. We next used our simulations to infer the ATP-dependence of 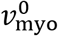 for fission yeast from the measurements by Mishra et al. (Mishra *et al*., 2013) of constriction rate versus ATP concentration (blue points, Fig. 7). The link between the two, provided by the simulations, is the dependence of constriction rate on 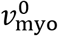 (Fig. 3j). Fitting to Michaelis-Menten kinetics yielded a maximal load-free velocity at saturating ATP of 0.23 μm s^−1^, close to the 0.24 μm s^−1^ reported in ref. (Stark *et al*., 2010), and a half-maximal load-free velocity at 30 μM ATP (Supplemental Note 2). These values are in the context of the cytokinetic ring machinery, and we note that 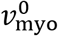 is a collective molecular property reflecting the complexities of the contractile ring architecture and interactions. For comparison, a half-maximal value 50 μM ATP was measured in vitro for skeletal muscle myosin (Kron and Spudich, 1986). Related in vitro measurements in fission yeast were performed in ref. (Pollard *et al*., 2017), where the ATpase rate of fission yeast myosin Myo2 was measured versus actin concentration at saturating ATP levels.

**Fig. 7.**
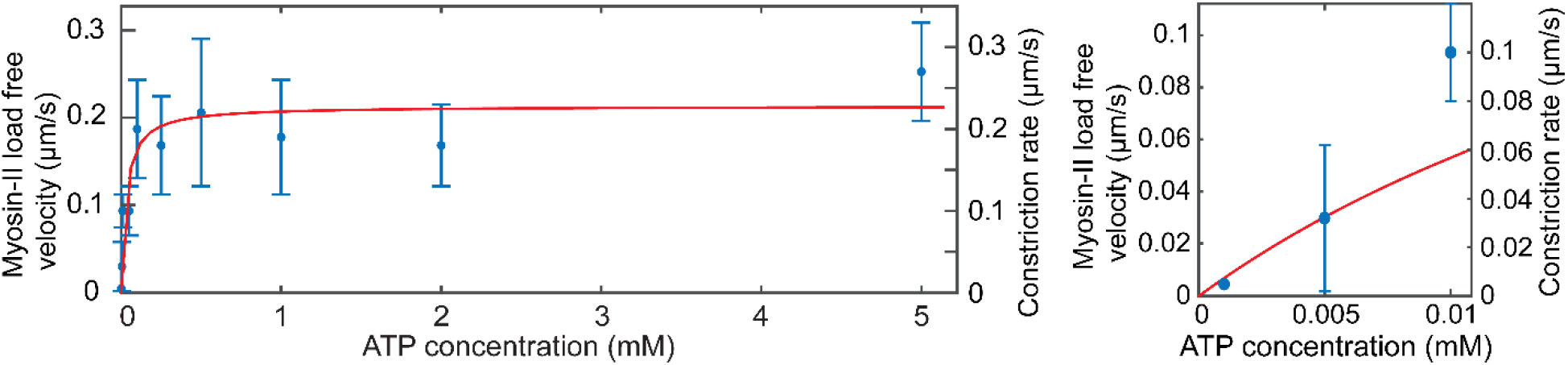
Myosin-II load free velocity and constriction rate versus ATP concentration. Constriction rates are experimental values from ref. (Mishra *et al*., 2013). The corresponding Myosin-II velocities (i.e., the scale for the vertical axis at left) were obtained from the experimental constriction rates using best fit line in Fig. 3j. Red curve: best fit Michaelis-Menten relation, corresponding to a maximal load-free velocity at saturating ATP of 0.23 μm s^−1^ and a half-maximal load-free velocity at 30 μM ATP (Supplementary Note 2). Plot at right: blow up near origin.

### Ring constriction in permeabilized protoplasts does not require actin turnover

It has been proposed that ring constriction is driven by actin depolymerization (Mendes Pinto *et al*., 2012). To test this, Mishra et al. used a cofilin mutant or the F-actin stabilizing drug jasplakinolide (Mishra *et al*., 2013). Constriction rates were unaffected. To mimic these experiments, we ran simulations with cofilin-mediated severing abolished or reduced. In agreement with experiment, constriction rates were unaltered (Fig. 6c,d).

This finding is as expected, because in the reeling-in mechanism the constriction rate is set by 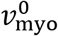, which is unaffected by the lengths of actin filaments in the ring. Hence no dependence of constriction rate on cofilin or other actin polymerization/depolymerization factors is expected. Thus, our model explains why actin turnover is not required for constriction of partially unanchored rings.

## Discussion

The cytokinetic ring plays center stage during cytokinesis, and its ability to generate tension and constrict is critical to cell division. How it produces tension remains unsettled, in part because many organizational details are unknown. Evidence for a muscle-like sarcomeric machinery is lacking, although some super-resolution microscopy and EM images show a degree of spatial periodicity of the organization in animal cells (Beach *et al*., 2014; Fenix *et al*., 2016; Henson *et al*., 2017). The ring organization appears to exhibit considerable randomness (Green *et al*., 2012; Laplante *et al*., 2015), so that the mechanism appears something of a puzzle given that a theoretical actomyosin bundle with randomly organized actin filaments and myosin-II is not tensile. Indeed, our simulations of free-standing randomly organized actomyosin rings produced no tension (Fig. 6).

Our analysis showed that the experiments of Mishra et al. (Mishra *et al*., 2013) provide quantitative evidence that fission yeast solves this problem simply by anchoring actin filaments at their barbed ends to the membrane in the ring. Conceptually, this is a natural way to create tension out of disorder, as every myosin-actin interaction renders the filament involved tensile (Fig. 6e). (Compare this with the random, unanchored bundle, where dynamically crosslinked myosin-propelled actin filaments are as often tensile as compressive.) The net ring tension is the sum effect of these tensile filament contributions without the need for a particular organization, sarcomeric or other.

Here we built a model implementing this organization, which quantitatively reproduced the observations of ref. (Mishra *et al*., 2013): (1) Unanchored ring segments had zero tension (Fig. 3a), (2) with no fitting parameters segments shortened at close to the zero-load velocity of myosin-II, with mean rate 0.24 μm s^−1^ very close to the experimental value 0.22 μm s^−1^ (Fig. 3), and (3) the shortening rate was independent of initial length (Fig. 3g). Anchoring schemes other than barbed-end anchoring could not explain these findings (Figs. 5c,d).

The experimental observations, (1)-(3), are inconsistent with a passive elastic mechanism, which would generate tension in an ananchored segment, or with a sarcomeric-like organization of interconnected contractile units, which would shorten at a rate proportional to the number of sarcomeric units and initial length. In *Caenorhabditis elegans* embryos, for example, shortening rates in successive divisions are proportional to initial ring length, not inconsistent with a sarcomeric-like organization (Carvalho *et al*., 2009). By contrast, length-independent shortening,(3), is a hallmark of the reeling-in mechanism we identified (Fig. 5), which acts at the ends of an unanchored segment unaffected by the intervening segment length.

Thus, both radial and lateral anchoring feature in the contractile ring machinery. Radial anchoring attaches the ring to the membrane and conveys the ring tension to the membrane, the cortex and, for fungi such as fission yeast, the cell wall. Lateral anchoring restrains lateral motion of ring components and features in the tension mechanism by retarding lateral sliding of actin filament barbed end anchors in the membrane, allowing the filaments to develop tension when pulled by myosin (Fig. 6e). The requirement for tension is that the lateral sliding velocity be well below 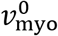, a condition that is satisfied in fission yeast (Laplante *et al*., 2016).

An implicit assumption of the present analysis is that the unanchoring process does not leave behind myosin-II and other core ring components in the membrane. In support of this assumption, detached ring segments remained intact and contained dynamic myosin-II Rlc1p, showing that some or all of the F-actin and myosin-II pulled away intact from the plasma membrane. Moreover, in the images of ref. (Mishra *et al*., 2013) myosin fluorescence on the membrane is not apparent in the vicinity of detached ring segments.

As a contractile cellular machine the cytokinetic ring presumably has a signature tension-constriction rate relationship, analogously to muscle (Edman, 1979) and other actomyosin systems. Most of this relationship is normally invisible outside a narrow physiological operating range, but yeast cell ghosts provide a laboratory to study the contractile ring in extraordinary circumstances outside this window. This can reveal otherwise hidden aspects of the workings of the machinery.

In fission yeast two extremes of behavior corresponding to two regions of the tension-constriction rate relation are now characterized. (1) In normal cells the ring constricts slowly compared to component turnover rates, operating near the high load isometric tension limit (Stachowiak *et al*., 2014; Thiyagarajan *et al*., 2015). The ring sets the tension to the isometric value, the value at fixed ring length and an intrinsic property of the ring (Stachowiak *et al*., 2014), and does not set the constriction rate. For example, in yeast protoplasts rings constrict along the membrane at various speeds depending on the surface steepness (Mishra *et al*., 2012; Stachowiak *et al*., 2014), showing that the constriction rate is not intrinsic to the ring. Indeed, in normal yeast cells the constriction rate is the rate of septum closure, and experiment and modeling suggest there is almost no influence from ring tension (Thiyagarajan *et al*., 2015). (2) Here we studied the opposite extreme in cell ghosts: fast, load-free constriction of partially anchored ring segments. In the load-free limit the tension vanishes, and the ring constricts via a novel reeling-in mechanism in which the ring itself sets the constriction rate. In this mode the constriction rate is indeed an intrinsic property of the ring, a multiple of the load-free myosin-II velocity, 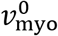.

## Materials and Methods

Methods and any associated references are described in Supplementary Note 1.

## ACKNOWLEDGEMENTS

We thank Sathish Thiyagarajan and Harvey F. Chin for helpful discussions. Research reported in this publication was supported by the National Institute Of General Medical Sciences of the National Institutes of Health under Award Number R01GM086731. The content is solely the responsibility of the authors and does not necessarily represent the official views of the National Institutes of Health.

## AUTHOR CONTRIBUTIONS

B. O’S. conceived the study, B.O’S. and S.W. designed the model, S.W. performed the simulation and B.O’S. and S.W analyzed the data. B.O’S. and S.W. wrote the manuscript.

The authors declare no competing financial interests.

## References

Arasada, R., and Pollard, T.D. (2014). Contractile ring stability in S. pombe depends on F-BAR Protein Cdc15p and Bgs1p transport from the Golgi complex. Cell Rep. 8, 1533–1544.

Beach, J.R., Shao, L., Remmert, K., Li, D., Betzig, E., and Hammer, J.A., 3rd. (2014). Nonmuscle myosin II isoforms coassemble in living cells. Curr. Biol. 24, 1160–1166.

Broersma, S. (1960). Viscous Force Constant for a Closed Cylinder. J. Chem. Phys. 32, 1632.

Carvalho, A., Desai, A., and Oegema, K. (2009). Structural memory in the contractile ring makes the duration of cytokinesis independent of cell size. Cell 137, 926–937.

Edman, K.A. (1979). The velocity of unloaded shortening and its relation to sarcomere length and isometric force in vertebrate muscle fibres. J. Physiol. 291, 143–159.

Fenix, A.M., Taneja, N., Buttler, C.A., Lewis, J., Van Engelenburg, S.B., Ohi, R., and Burnette, D.T. (2016). Expansion and concatenation of non-muscle myosin IIA filaments drive cellular contractile system formation during interphase and mitosis. Mol. Biol. Cell.

Fujiwara, K., and Pollard, T.D. (1976). Fluorescent antibody localization of myosin in the cytoplasm, cleavage furrow, and mitotic spindle of human cells. J. Cell Biol. 71, 848–875.

Goss, J.W., Kim, S., Bledsoe, H., and Pollard, T.D. (2014). Characterization of the roles of Blt1p in fission yeast cytokinesis. Mol. Biol. Cell 25, 1946–1957.

Green, R.A., Paluch, E., and Oegema, K. (2012). Cytokinesis in animal cells. Annu. Rev. Cell Dev. Biol. 28, 29–58.

Henson, J.H., Ditzler, C.E., Germain, A., Irwin, P.M., Vogt, E.T., Yang, S., Wu, X., and Shuster, C.B. (2017). The ultrastructural organization of actin and myosin II filaments in the contractile ring: new support for an old model of cytokinesis. Mol. Biol. Cell 28, 613–623.

Hiramoto, Y. (1975). Force exerted by cleavage furrow of sea-urchin eggs. Dev. Growth Differ. 17, 27–38.

Kielty, C.M., Sherratt, M.J., and Shuttleworth, C.A. (2002). Elastic fibres. J. Cell Sci. 115, 2817–2828.

Kovar, D.R., Harris, E.S., Mahaffy, R., Higgs, H.N., and Pollard, T.D. (2006). Control of the assembly of ATP- and ADP-actin by formins and profilin. Cell 124, 423–435.

Kron, S.J., and Spudich, J.A. (1986). Fluorescent actin filaments move on myosin fixed to a glass surface. Proc. Natl. Acad. Sci. USA 83, 6272–6276.

Laplante, C., Berro, J., Karatekin, E., Hernandez-Leyva, A., Lee, R., and Pollard, T.D. (2015). Three Myosins Contribute Uniquely to the Assembly and Constriction of the Fission Yeast Cytokinetic Contractile Ring. Curr. Biol. 25, 1955–1965.

Laplante, C., Huang, F., Tebbs, I.R., Bewersdorf, J., and Pollard, T.D. (2016). Molecular organization of cytokinesis nodes and contractile rings by super-resolution fluorescence microscopy of live fission yeast. Proc. Natl. Acad. Sci. U. S. A. 113, E5876–E5885.

Mabuchi, I., and Okuno, M. (1977). The effect of myosin antibody on the division of starfish blastomeres. J. Cell. Biol. 74, 251–263.

Mendes Pinto, I., Rubinstein, B., Kucharavy, A., Unruh, J.R., and Li, R. (2012). Actin depolymerization drives actomyosin ring contraction during budding yeast cytokinesis. Dev. Cell 22, 1247–1260.

Mishra, M., Huang, Y., Srivastava, P., Srinivasan, R., Sevugan, M., Shlomovitz, R., Gov, N., Rao, M., and Balasubramanian, M. (2012). Cylindrical cellular geometry ensures fidelity of division site placement in fission yeast. J. Cell Sci. 125, 3850–3857.

Mishra, M., Kashiwazaki, J., Takagi, T., Srinivasan, R., Huang, Y., Balasubramanian, M.K., and Mabuchi, I. (2013). In vitro contraction of cytokinetic ring depends on myosin II but not on actin dynamics. Nat. Cell Biol. 15, 853–859.

Naqvi, N.I., Eng, K., Gould, K.L., and Balasubramanian, M.K. (1999). Evidence for F-actin-dependent and -independent mechanisms involved in assembly and stability of the medial actomyosin ring in fission yeast. EMBO J. 18, 854–862.

O’Shaughnessy, B., and Thiyagarajan, S. (2018). Mechanisms of contractile ring tension production and constriction. Biophys Rev 10, 1667–1681.

Pelham, R.J., and Chang, F. (2002). Actin dynamics in the contractile ring during cytokinesis in fission yeast. Natur 419, 82–86.

Pollard, L.W., Bookwalter, C.S., Tang, Q., Krementsova, E.B., Trybus, K.M., and Lowey, S. (2017). Fission yeast myosin Myo2 is down-regulated in actin affinity by light chain phosphorylation. Proceedings of the National Academy of Sciences of the United States of America 114, E7236–E7244.

Pollard, T.D., and O’Shaughnessy, B. (2019). Molecular Mechanism of Cytokinesis. Annu. Rev. Biochem. 88, null.

Rappaport, R. (1967). Cell division: direct measurement of maximum tension exerted by furrow of echindoerm eggs. Science 156, 1241–1243.

Rappaport, R. (1977). Tensiometric studies of cytokinesis in cleaving sand dollar eggs. J. Exp. Zool. 201, 375–378.

Schroeder, T.E. (1972). The contractile ring. II. Determining its brief existence, volumetric changes, and vital role in cleaving Arbacia eggs. J. Cell Biol. 53, 419–434.

Schroeder, T.E., and Otto, J.J. (1988). Immunofluorescent Analysis of Actin and Myosin in Isolated Contractile Rings of Sea-Urchin Eggs. Zool. Sci. 5, 713–725.

Stachowiak, M.R., Laplante, C., Chin, H.F., Guirao, B., Karatekin, E., Pollard, T.D., and O’Shaughnessy, B. (2014). Mechanism of cytokinetic contractile ring constriction in fission yeast. Dev. Cell 29, 547–561.

Stark, B.C., Sladewski, T.E., Pollard, L.W., and Lord, M. (2010). Tropomyosin and myosin-II cellular levels promote actomyosin ring assembly in fission yeast. Mol. Biol. Cell 21, 989–1000.

Thiyagarajan, S., Munteanu, E.L., Arasada, R., Pollard, T.D., and O’Shaughnessy, B. (2015). The fission yeast cytokinetic contractile ring regulates septum shape and closure. J. Cell Sci.

Vavylonis, D., Wu, J.Q., Hao, S., O’Shaughnessy, B., and Pollard, T.D. (2008). Assembly mechanism of the contractile ring for cytokinesis by fission yeast. Sci 319, 97–100.

Wu, J.Q., and Pollard, T.D. (2005). Counting cytokinesis proteins globally and locally in fission yeast. Sci 310, 310–314.

Yoneda, M., and Dan, K. (1972). Tension at the surface of the dividing sea-urchin egg. J. Exp. Biol. 57, 575–587.

Yonetani, A., Lustig, R.J., Moseley, J.B., Takeda, T., Goode, B.L., and Chang, F. (2008). Regulation and targeting of the fission yeast formin cdc12p in cytokinesis. Mol. Biol. Cell 19, 2208–2219.

