## Supplementary Material for "Anchoring of actin to the plasma membrane enables tension production in the fission yeast cytokinetic ring"

### Supplementary Note 1
### Model of the Partially Unanchored Fission Yeast Cytokinetic Ring

#### Contents


1. **3D molecularly explicit model of the cytokinetic ring in permeabilized fission yeast protoplasts**
   a. **Introduction**
   b. **Anchored ring segment**
   c. **Unanchored ring segment**
   d. **Interfaces between anchored and unanchored segments**
2. **Simulation of the model: constriction of partially unanchored rings**
   a. **Initial condition: the steady state cytokinetic ring in normal, intact cells**
   b. **Computational scheme**
   c. **Calculation of ring length**
   d. **Calculation of ring constriction rate**
   e. **Calculation of ring tension**
   f. **Calculation of ring component densities**
3. **Determination of model parameters**
   a. **Determination of myosin-II load-free velocity $v^0_{\mathrm{myo}}$ from motility assay of ref. (1)**
   b. **Turnover parameters**
4. **The reeling-in constriction mechanism**

### 1. 3D molecularly explicit model of the cytokinetic ring in permeabilized fission yeast protoplasts

#### a. Introduction

Here we present the model and computer simulation scheme we developed to describe the constriction kinetics of cytokinetic rings measured by Mishra et al. (2). In these experiments, fission yeast protoplasts were prepared by treating normal yeast cells to remove the cell wall, and the protoplast membranes were permeabilized with a detergent, causing the loss of cytoplasm and cytoplasmic structures. The resultant cell ghosts provided a laboratory for the authors to study the cytokinetic ring: the medium permeating the cells can be controlled, and rings are subject to extraordinary circumstances in which otherwise hidden behaviors occur that can reveal fundamental information about the ring constriction mechanism.

A quantitative model of ring constriction in permeabilized protoplasts must describe partially unanchored rings. Mishra et al. found that, when ATP was added to the medium to trigger contractility in cells with cytokinetic rings, rings constricted at a much higher rate than in normal cells (2). In all reported time lapse images of constricting rings, the mode of constriction was as follows: first, one or more segments of the ring appeared to detach from the plasma membrane, i.e. became unanchored; subsequently, the unanchored segment shortened until it became straight and apparently taut. The shortening rate was ~30-fold faster than in normal cells, was independent of the initial length of the ring or the unanchored segment, and was constant in time. Anchored segments in the same ring did not shorten.

Some critical aspects of the model we developed are described below.

**(i) <u>The model describes partially anchored rings.</u>** To mirror the experiments of Mishra et al., we applied the model to the situation when individual rings have both anchored and unanchored sections. One segment of the ring is unanchored from the plasma membrane, while the remainder is normally anchored (2) (see Figs. 1, 2b of main text). In the unanchored segment the anchors of all components are removed from the membrane; thus, component motions are unaffected by drag forces from the plasma membrane that severely retard the motions of anchored components, and the components are not constrained to move within the surface of the membrane. Instead, components move through the aqueous medium with velocities set by the forces acting upon them and very small viscous drag forces due to the aqueous (non-cytosolic) medium (Fig. 2b of main text). An essential feature of the model is the interface between the anchored and unanchored segments, which turns out to play a critical role in the mechanism of constriction (Fig. 5 of main text).

**(ii) <u>The model is molecularly explicit.</u>** As the fission yeast cytokinetic ring is uniquely well characterized, it offers the best opportunity to construct a realistic mathematical model with minimal assumptions (3). Over 150 gene products have been identified (4), the biochemical properties of many key components have been characterized, and the amounts of more than 25 contractile ring proteins were measured as a function of time throughout the course of constriction (5-7).

The model is coarse-grained by design, to capture the collective behavior of the thousands of molecular components in the ring. However, the key components are explicitly represented, using the large body of available experimental information to characterize these components (Table S1 shows the key parameter values used in the model and the experimental sources; all other parameters are specified in the main text). For example, F-actin filaments are represented as rods that can assume any 3D shape, determined by the forces exerted upon them and the bending stiffness of F-actin which we take from experiment. The actin filaments are anchored by formin proteins to the plasma membrane, with an anchor drag coefficient determined from experiment (see Fig. 2a of main text). The formins nucleate and polymerize filaments in arbitrary directions, and the filaments are subject to stochastic severing by ADF-cofilin (see Fig. 2a of main text). The polymerization and severing rates are determined from experimental data.

At the instant of ATP addition, the simulated ring contains 180-285 actin filaments depending on the initial ring length. This range of values is based on the measured number of formin Cdc12p dimers in normal cells (7), assumed to equal the number of actin filaments, and the assumption that the formin density per length of ring in protoplasts and normal cells are equal. Each actin filament is represented by a series of subunits, each subunit being a bead. The subunits are connected by rods of length 100 nm. Myosin-II clusters have an effective size ~100 nm (3, 8), the capture radius for myosin-actin binding, and the model does not describe smaller scale details within the clusters.

**(iii) The model is 3-dimensional.** In the model the contour of the ring can follow any closed curve in 3D space, and the detailed structure of the ring (i.e. positions and orientations of ring components) is described across its width, its thickness and its length. Actin filaments, for example, are anchored at their barbed ends to the plasma membrane by formin dimers and orient in arbitrary directions away from the membrane (see Fig. 2a of main text). Component motions are tracked in all directions as, for example, when a segment of the ring detaches from the membrane and is pulled through the aqueous medium (2) (Figs. 1, 2b). In this case the components of the ring experience forces that tend to pull them away from the membrane, while viscous drag forces from the surrounding medium oppose this motion.

**(iv) The model is fully dynamic.** In addition to generating tension in the ring and evolving the component locations and configurations, the model constricts the ring: the length of the ring is directly determined by the motions of the ring components and is continuously updated as the simulation proceeds. The force-velocity relation for myosin-II is incorporated (see Sec. 1b below), and the model can describe fast component motions and high constriction rates. This is needed to describe constriction in permeabilized protoplasts where constriction rates were 30-fold the normal rate (2).

**(v) The model describes the particular turnover kinetics of rings in permeabilized protoplasts.** In permeabilized protoplasts most cytosolic components are absent. Thus, we exclude binding of new components that normally replenish dissociating components (3, 9, 10) (see Fig. 2 of main text). Further, the absence of cytosolic components apparently also affected dissociation rates: from measurements by Mishra et al. (2) of the amount of myosin-II regulatory light chain Rlc1p and the amount of actin remaining in the ring after constriction, we deduced the dissociation rate constants for these and other components. These rate constants were lower than in normal cells (see Table S1 and Sec. 3b below).

Details of the model follow. The ring components and their interactions are similar to those in our previous model of the normal fission yeast cytokinetic ring, the Stachowiak model (3). The Stachowiak model describes a fully anchored ring, is a 2D representation (i.e. the ring is a ribbon attached to the plasma membrane) and the ring length $L$ is fixed, i.e. the model is not fully dynamic. In the present analysis of constriction in permeabilized protoplasts, we assume the cytokinetic ring has previously attained a normal steady state prior to permeabilization, i.e. the ring has been frozen in this state until the instant of ATP addition that triggers contraction and partial detachment of the ring. To describe this steady state, which serves as an initial condition for the present analysis, we used the results of the Stachowiak model of the fully anchored ring to fix the configurations of all ring components (Sec. 2a below).

#### b. Anchored ring segment

**Evidence that components in the normal ring are anchored.** Previous measurements suggest that formin Cdc12 and myosin-II in the ring are anchored to the plasma membrane. We previously measured motions of fluorescent spots that contained formin Cdc12p and myosin-II regulatory light chain Rlc1p in constricting fission yeast protoplast cytokinetic rings (3). From kymographs we found that both formin and myosin-II moved at speeds much less than the myosin-II load-free velocity, suggesting that both components are anchored to the plasma membrane and that myosin-II works against large anchor drag forces. We used simulations of the Stachowiak model together with the experimentally measured velocities to deduce the anchor drag coefficients of each component (3).

A second suggestive fact is that formin and myosin-II are anchored to the plasma membrane in the precursor nodes from which the ring is assembled (11). Third, myosin-II remains at the division site after disassembly of actin filaments in the fission yeast ring (12), suggesting that myosin-II may be anchored to the plasma membrane.

In the present study we assume that in normal segments of the ring (those which have not detached) formin dimers and myosin-II clusters are anchored to the plasma membrane, with anchor drag coefficients based on the best-fit values obtained in ref. (3) (Table S1).

#### (i) Ring components

*Formin-capped actin filaments.* The model treats actin filaments as semi-flexible, with a bending modulus $\kappa = k_B T l_p$, where $k_B$ is Boltzmann's constant, $T$ is the temperature and $l_p$ = 10 μm the persistence length (13). The filaments are assumed inextensible, a justified approximation because the extension of an actin filament under physiological conditions is negligible. Taking a typical value of 400 pN for the ring tension (3), distributed among ∼20 actin filaments (14) in the cross-section of the ring gives an average of ∼20 pN tension per actin filament. A stiffness of 65.3 pN/nm was reported for a 1-μm-long actin filament (15), giving a stiffness 50.2 pN/nm for a filament with the average length in the ring of 1.3 μm (3). This gives a negligible extension of 0.4 nm.

We track every 37[th] actin subunit along the filament, corresponding to a distance $l_0 = 0.1$ μm along the filament (3).Thus each actin filament is represented as a series of subunits connected by rigid rods. A membrane-anchored formin dimer caps the barbed end of every actin filament.

*Myosin-II clusters.* Myosin-II clusters are anchored to the plasma membrane. Since the precise organization of myosin-II in the fission yeast cytokinetic ring during constriction is not established, we assume an organization with clusters of uniform size, each with 8 myosin dimers. The Stachowiak model (3) assumed 20 dimers per cluster, the reported (7) number of myosin-II dimers in the 'nodes' (protein complexes) from which the fission yeast ring is assembled. We found that 20 dimers per cluster resulted in catastrophic fracture of the ring; since no such events were reported by Mishra et al. (2), we used a smaller number that ensured stability.

*Crosslinking by α-actinin.* Actin filaments are crosslinked by α-actinin dimers. We model each α-actinin crosslink as a spring connecting two actin subunits on different filaments (3), with spring constant $k_x = 25$ pN/μm and rest length $r_x^0 = 30$ nm. The crosslinks are dynamic, with an intrinsic off rate $k_{off}^x \sim 3.3\ s^{-1}$ (3, 16-18). Further, we assume that two subunits become uncrosslinked if their separation exceeds the maximum length of the crosslink; thus, the crosslinks dissociate if the separation of the linked actin subunits exceeds $r_x^{bind} = 50$ nm.

Simulations of the model with α-actinin showed that only $1.0 \pm 0.5\%$ of the α-actinin crosslinks initially present were still present after 1 s, and virtually all crosslinks had dissociated within ~ 2 s. This was due to the large intrinsic off rate, and over-stretching by moving actin filaments. This time scale ~ 1 s is much smaller than the ring constriction time (~ 25-60 s); accordingly, we found that simulated constriction rates were unaffected when α-actinin was altogether omitted from the simulations (see Fig. S2). Moreover, in our previous study we showed that in normal cells α-actinin contributes $< 1\%$ of ring tension (3). For these reasons, all other simulations in our study were run without α-actinin to minimize simulation running times.

#### (ii) Forces

*Myosin-II capture.* In the model a myosin-II cluster binds to any actin subunit within a certain capture radius $r_{\text{myo}}$ and draws the subunit towards the center of the cluster. This binding interaction is implemented as a spring that has zero rest length and connects the center of the myosin-II cluster to the actin subunit. To avoid adding to or subtracting from the pulling force of the myosin, the component of the capture force perpendicular to the actin filament is used:

$$\boldsymbol{f}_{i,\alpha}^{\text{cap}} = -k_{\text{myo}} \left\{(\boldsymbol{r_i} - \boldsymbol{r_\alpha}) - [(\boldsymbol{r_i} - \boldsymbol{r_\alpha}) \cdot \widehat{\mathbf{T}}_i]\widehat{\mathbf{T}}_i\right\} \quad (1)$$

where $\boldsymbol{f}_{i,\alpha}^{\text{cap}}$ is the capture force exerted on actin subunit $i$ by myosin-II cluster $\alpha$, $\boldsymbol{r_i}$ and $\boldsymbol{r_\alpha}$ are their positions, and $k_{\text{myo}} = 40\ \text{pN}/\mu\text{m}$ is the equivalent spring constant. The first term in the curly brackets is the position vector from the cluster to the subunit, and the second term subtracts off from the first term the component parallel to the filament. $\widehat{\mathbf{T}}_i$ is the unit tangent vector of the actin filament at subunit $i$, discretized as

$$\widehat{\mathbf{T}}_i = \begin{cases} \dfrac{\boldsymbol{r}_{i+1} - \boldsymbol{r}_i}{|r_{i+1} - r_i|}, \text{if subunit } i \text{ is not at the pointed end} \\ \dfrac{\boldsymbol{r}_i - \boldsymbol{r}_{i-1}}{|r_i - r_{i-1}|}, \text{if subunit } i \text{ is at the pointed end} \end{cases} \quad (\mathbf{2})$$

If more than one subunit on the same filament is within $r_{\text{myo}}$ of a myosin-II cluster, the force is exerted only on the subunit closest to the pointed end. The force on the myosin-II cluster $\alpha$ exderted by the actin subunit $i$ is $-\boldsymbol{f}_{i,\alpha}^{\text{cap}}$.

*Myosin-II pulling force.* A myosin-II cluster pulls every actin subunit within its capture radius $r_{\text{myo}}$ with a force tangent to the filament (see Fig. 2a of main text). We use a linear force-velocity relation: the pulling force $\boldsymbol{f}^{\text{pull}}$ decreases linearly with the speed that the myosin-II cluster moves along the filament. If more than one subunit on the same filament is within $r_{\text{myo}}$ of a myosin-II cluster, the force is exerted only on one subunit, to avoid overcounting; the selected subunit is that which is furthest from the barbed end, unless that subunit is the pointed end. This ensures numerical stability. For such an interacting myosin cluster/actin subunit pair, the pulling force on the actin subunit $i$ by the myosin-II cluster α is given by

$$\boldsymbol{f}_{i,\alpha}^{\text{pull}} = f_{\text{s}} \left[1 - \frac{(\boldsymbol{v_i} - \boldsymbol{v_\alpha}) \cdot \hat{\mathbf{t}}_i}{v_{\text{myo}}^0}\right] \hat{\mathbf{t}}_i \quad (\mathbf{3})$$

where $\boldsymbol{v_i}$ and $\boldsymbol{v_\alpha}$ are the velocities of the actin subunit and myosin-II cluster, $f_{\text{s}}$ is the myosin-II stall force per cluster, and $\hat{\boldsymbol{t}}_i = (\boldsymbol{r_{i+1}} - \boldsymbol{r_i})/|\boldsymbol{r_{i+1}} - \boldsymbol{r_i}|$ is the unit vector pointing from the ($i$-1)$^{\text{th}}$ subunit to the $i^{\text{th}}$ subunit. The pulling force on the myosin-II cluster $\alpha$ is $-\boldsymbol{f}_{i,\alpha}^{\text{pull}}$.

*Myosin-II saturation effects.* We take a stall force per myosin-II cluster of $f_{\text{s}} = 4$ pN from our previous experimental measurements of node motions in protoplasts (3) and in intact cells (11). The meaning of this stall force is the force (at zero velocity) exerted by one myosin-II cluster on one filament that it interacts with. It follows that the total force exerted by a cluster that interacts with $n_{\text{fil}}$ filaments is equal to (4 pN) $n_{\text{fil}}$. Now, if we assume a stall force of 2 pN per myosin-II *dimer* (19), the maximum total force a cluster could exert upon all filaments it interacts with is 16 pN, since we assume 8 dimers per cluster. For this reason, in the case that $n_{\text{fil}} > 4$, the stall force is lowered to a value $f_{\text{s}} = (4\ \text{pN}) \times (4/n_{\text{fil}})$ per cluster per actin filament the cluster interacts with. In other words, we assume that a myosin-II cluster is *saturated* when 4 or more actin filaments interact with it, and we ensure that the total force summed over all filaments the cluster interacts with is equal to the maximum allowed value of 16 pN. This means that for 5 or more filaments the (stall) force exerted per filament is lowered. Our algorithm distributes this total force *evenly* among the $n_{\text{fil}}$ filaments.

*Myosin-II excluded volume.* Myosin-II clusters repel one other if they move to within a distance $d_{\text{myo}} = 40$ nm of each other. For two clusters $\alpha$ and $\beta$ within $d_{\text{myo}}$, the repulsive force on $\alpha$ is

$$\boldsymbol{f}_{\alpha,\beta}^{\text{excl}} = -k_{\text{myo}}^{\text{excl}}\left(d_{\text{myo}} - |\boldsymbol{r}_{\beta} - \boldsymbol{r}_{\alpha}|\right)\frac{\boldsymbol{r}_{\beta} - \boldsymbol{r}_{\alpha}}{|\boldsymbol{r}_{\beta} - \boldsymbol{r}_{\alpha}|} \quad (4)$$

while the repulsive force on $\beta$ is $-f_{\alpha,\beta}^{\text{excl}}$. Here the elastic constant $-k_{\text{myo}}^{\text{excl}} = 4$ pN/nm. The total repulsive force on $\alpha$ due to all clusters $\{\beta||\boldsymbol{r}_{\beta} - \boldsymbol{r}_{\alpha}| < d_{myo}\}$ is

$$\boldsymbol{f}_{\alpha}^{\text{excl}} = \sum_{\{\beta||\boldsymbol{r}_{\beta}-\boldsymbol{r}_{\alpha}|<d_{\text{myo}}\}} \boldsymbol{f}_{\alpha,\beta}^{\text{excl}} \quad (5)$$

*Tension in individual actin filaments.* The tension that an actin filament bears is represented by the pairwise attractive forces that act between adjacent actin subunits. Between subunits $i$ and $i+1$, the force on the subunit $i$ is $f_{i,i+1}^{\text{tension}}(\boldsymbol{r}_{i+1} - \boldsymbol{r}_{i})/|\boldsymbol{r}_{i+1} - \boldsymbol{r}_{i}|$, and the force on the subunit $i+1$ is minus this value. The magnitude of this force, $f_{i,i+1}^{\text{tension}}$, can be thought of as the tension in the rigid rod that connects the two subunits and is calculated together with the velocities of the ring components at every time step (see section "Computation scheme" below).

*Actin filament bending force.* The discretized version of the bending energy of a filament with bending modulus $\kappa$ is (20)

$$H^{B} = \frac{\kappa}{l_0}\sum_{i=2}^{N-1}(1 - \hat{\boldsymbol{t}}_{i} \cdot \hat{\boldsymbol{t}}_{i-1}) \quad (6)$$

where $l_0 = 0.1$ μm is the separation between adjacent actin subunits on a filament and $N$ is the total number of subunits on the filament. The bending force on each subunit is calculated as minus the derivative of $H^{B}$ with respect to the coordinates of the subunit.

*α-actinin crosslinking.* An α-actinin crosslink that connects actin subunits $i$ and $j$ is represented as a spring, exerting elastic forces $\boldsymbol{f}_{i,j}^{\text{x}} = -k_{\text{x}}\left(|\boldsymbol{r}_{i} - \boldsymbol{r}_{j}| - r_{\text{x}}^{0}\right)(\boldsymbol{r}_{i} - \boldsymbol{r}_{j})/|\boldsymbol{r}_{i} - \boldsymbol{r}_{j}|$ on subunit $i$ and $-\boldsymbol{f}_{i,j}^{\text{x}}$ on subunit $j$. Therefore, the total crosslinking force on subunit $i$ is

$$\boldsymbol{f}_{i}^{\text{x}} = \sum_{\{j|i \text{ and } j \text{ linked}\}} \boldsymbol{f}_{i,j}^{\text{x}} \quad (7)$$

*Confinement of ring components by the membrane.* Components of the ring cannot pass through the plasma membrane. Even in the experiments of Mishra et al. (2) where the membranes were permeabilized, ring components did not appear to escape the volume enclosed by the plasma membrane of the cell. To impose this constraint, we include in our model and artificial elastic restoring force $\boldsymbol{f}^{\text{mb}} = -k_{\text{mb}}\left(\sqrt{x^2 + y^2 + z^2} - R\right)\hat{r}$ acting on every formin, myosin-II cluster and actin subunit that stray outside of the volume enclosed by the membrane, a sphere of radius $R$. The force is zero on every ring component within the volume enclosed by the membrane. Here $k_{\text{mb}} = 20$ pN/μm is the elastic constant and $\hat{r} = (\hat{x} + \hat{y} + \hat{z})/\sqrt{3}$ is the unit radial vector. The value of $k_{\text{mb}}$ is chosen to be strong enough to prevent significant unphysical detours by ring components, while not being so large as to force use of a very small simulation time step.

*Constraining component anchors to lie on the plasma membrane surface: normal anchoring force.* In the anchored segment of the ring, formin Cdc12 dimers and myosin-II clusters are anchored to the plasma membrane (3). To constrain their anchors to lie on the surface of the membrane, we use an anchoring force $\boldsymbol{f}^{\text{anch}} = f^{\text{anch}}\,\hat{r}$ that acts perpendicular to the plasma membrane (not to be confused with $\boldsymbol{f}^{\mathbf{mb}}$, see above). At every time step, the magnitude of this force, $f^{\mathrm{anch}}$, is recalculated together with the velocities of the ring components to ensure that the components anchors remain on the membrane surface (see section "Computation scheme").

*Membrane drag forces: tangential anchoring forces.* Formin Cdc12 dimers and myosin-II clusters that are anchored to the membrane are subject to membrane drag forces $\boldsymbol{f}_{\mathbf{for}}^{\mathbf{drag,mb}} = -\gamma_{\mathrm{for}}\boldsymbol{v}$ and $\boldsymbol{f}_{\mathbf{myo}}^{\mathbf{drag,mb}} = -\gamma_{\mathrm{myo}}\boldsymbol{v}$, respectively, when they move within the plasma membrane surface (Fig. 2a of main text). Here $\gamma_{\mathrm{for}}$ and $\gamma_{\mathrm{myo}}$ are the drag coefficients.

*Drag forces in aqueous medium.* In the experiments of Mishra et al., the cytokinetic ring constricted in the absence of most cytoplasmic constituents (2). The ring constricted through an aqueous solution with a viscosity presumably close to that of water. Our models uses an aqueous viscosity 5 times that of water, $\eta = 5 \times 10^{-3}$ Pa $\cdot$ s, to allow use of longer simulation time steps. We tested that ring constriction rates and ring tension were unaffected when the actual viscosity of water was used instead (i.e. 5-fold smaller drag coefficients), Fig. S3.

To calculate the drag coefficient $\gamma_{\mathrm{act}}$ of an actin subunit in the solution, we approximate it as a cylinder of length $l_0$ (the distance between adjacent actin subunits) and diameter $d_{\mathrm{act}} = 7$ nm, the diameter of an actin filament (21). The drag coefficients for such a cylinder in a fluid of viscosity $\eta$, for sideways and lengthwise motion, are $8\pi\eta l_0/(\sigma - g)$ and $4\pi\eta l_0/(\sigma - g)$, respectively, where $\sigma = \log(2l_0/d_{\mathrm{act}})$ and $g = 0.35 - 4(1/\sigma - 0.43)^2$ is a correction (22). Since there are significant sideways and lengthwise motions of the actin filaments in the simulation, we take an isotropic drag coefficient $\gamma_{\mathrm{act}} = 6\pi\eta l_0/\sigma \sim 3 \times 10^{-3}$ pN $\cdot$ s/μm for simplicity, the average of the sideways and lengthwise drag coefficients with the small correction $g$ omitted. The viscous force $\boldsymbol{f}_{\mathbf{act}}^{\mathbf{drag,bulk}}$ on an actin subunit is proportional to its velocity $\boldsymbol{v}$, $\boldsymbol{f}_{\mathbf{act}}^{\mathbf{drag,bulk}} = -\gamma_{\mathrm{act}}\boldsymbol{v}$.

##### (iii) Component dissociation

In the model formin, myosin-II and $\alpha$-actinin dissociate from the ring (Fig. 2a of main text). Actin dissociates through stochastic cofilin-mediated filament severing, and also via whole-filament unbinding together with a formin when the formin dissociates. Dissociating components are not replenished, being absent from the cytoplasm of permeabilized cells (2).

*Unbinding of formin dimers and myosin-II clusters from the ring.* After each time step, $n_{\mathrm{off}}^{\mathrm{for}} = k_{\mathrm{off}}^{\mathrm{for}}\Delta t$ formin dimers, together with the actin filaments that they bind, and $n_{\mathrm{off}}^{\mathrm{myo}} = k_{\mathrm{off}}^{\mathrm{myo}}\Delta t$ myosin-II clusters are deleted from the simulation, where $k_{\mathrm{off}}^{\mathrm{for}}$ and $k_{\mathrm{off}}^{\mathrm{myo}}$ are the formin and myosin off rates. The components selected for deletion are chosen randomly. In general, $n_{\mathrm{off}}^{\mathrm{for}}$ and $n_{\mathrm{off}}^{\mathrm{myo}}$ are not integers, and the actual numbers of formin dimers and myosin-II clusters deleted from the simulation were taken to be either the integer parts of $n_{\mathrm{off}}^{\mathrm{for}}$ and $n_{\mathrm{off}}^{\mathrm{myo}}$, or the integer parts plus unity, so as to yield mean values equal of $n_{\mathrm{off}}^{\mathrm{for}}$ and $n_{\mathrm{off}}^{\mathrm{myo}}$. We used this rule, based on an assumed integer number of dissociated components per time step, throughout this study.

*Cofilin-mediated severing of actin filaments.* We assume that every short segment of length $\Delta l$ on belonging to an actin filament has equal chance $P_{\mathrm{sev}} = r_{\mathrm{sev}}\,\Delta l\,\Delta t$ to be severed between time $t$ and $t + \Delta t$, where $r_{\mathrm{sev}}$ is the severing rate per filament length (3). At every time step, $n = r_{\mathrm{sev}} l_{\mathrm{tot}}\,\Delta t$ severing locations are selected at random, and all actin subunits between these points and the pointed ends are removed from the simulation.

*α-actinin dissociation.* At every time step, $n = k_{\text{off}}^{\text{x}} \Delta t$ α-actinin randomly selected crosslinks are deleted from the simulation, in addition to those that are over-stretched (see the section "α-actinin crosslinks" in "Ring components").

##### c. Unanchored ring segment

The unanchored segment features the same ring components as in the anchored segment, and most of the interactions are identical (Fig. 2b). The motions of components in the anchored segment, and the evolution of the unanchored ring segment as a whole, turn out of course to be completely different to those in the anchored segment. However, the major difference as far as the model is concerned is only that in the unanchored segment formin dimers and myosin-II clusters are not subject to anchor-membrane drag forces or anchoring forces normal to the membrane. Instead, they experience drag forces $\boldsymbol{f}_{\mathbf{for}}^{\mathbf{drag,bulk}} = -\gamma_{\text{for,bulk}} \boldsymbol{v}$ and $\boldsymbol{f}_{\mathbf{myo}}^{\mathbf{drag,bulk}} = -\gamma_{\text{myo,bulk}} \boldsymbol{v}$ from the aqueous solution. The drag coefficient of formin dimers is $\gamma_{\text{for,bulk}} = \gamma_{\text{act}}$ for simplicity. The drag coefficient for myosin-II clusters $\gamma_{\text{myo,bulk}}$ is calculated by approximating the myosin-II cluster as a sphere of radius $r_{\text{myo}}$, the capture radius. Its drag coefficient is given by the Stokes' Law expression, $\gamma_{\text{myo,bulk}} = 6\pi\eta r_{\text{myo}} \sim 0.01 \text{ pN} \cdot \text{s}/\mu\text{m}$.

##### d. Interfaces between anchored and unanchored segments

In our simulation there are two interfaces between the anchored segment and the unanchored segment of a ring (Fig. 2b). On one side of the interface, all formin dimers and myosin-II clusters are anchored, while on the other side all are unanchored. Some of the actin filaments anchored on the anchored side of the interface extend across the interface and into the unanchored side; these are the filaments which happen to be oriented in the direction of the unanchored segment. Similarly, some of the filaments connected to unanchored formin dimers on the unanchored side extend across the interface to the anchored side. During constriction of the ring, some of the formin dimers and myosin-II clusters will move across the interface. We assume that these components retain their anchoring status (anchored or unanchored) even after crossing the interface. Sec. 4 below presents a more detailed description of the interfaces, and how the interfaces drive ring constriction through a reeling-in mechanism.

#### 2. Simulation of the model: constriction of partially unanchored rings

##### a. Initial condition: the steady state cytokinetic ring in normal, intact cells

In the experiments of Mishra et al., cytokinetic rings appeared normal before the plasma membrane of the protoplast cells was permeabilized (2). Thus, we assume that prior to permeabilization the rings are normal rings characteristic of normal intact fission yeast cells, and we used our previously developed simulation of the fully anchored fission yeast ring to represent these rings, the Stachowiak model (3). This model is a molecularly explicit 2D representation. We used the statistical properties of the steady state ring generated by the Stachowiak model to set the initial condition of the present simulation, i.e to set the state of the ring at the instant prior to detachment of a ring segment.

The initial state of the ring is as follows. The ring is a 0.2 μm wide bundle of actin filaments and myosin-II clusters, with all actin filaments lying parallel to the bundle with a randomly clockwise or anti-clockwise orientation, anchored to the membrane at their barbed ends via formins that are randomly positioned along the ring (3). The actin filaments are slightly bent to follow the curvature of the membrane. Myosin-II clusters are anchored at random locations along the ring, independently of the formins and of one another, except that a minimum initial separation of $d_{\text{myo}}$ between any two myosin-II clusters is enforced. Both formin dimers and myosin-II clusters are uniformly distributed across the width of the ring. The length of the ring varies over the range 12-19 μm (2).

Initially, the actin filaments follow the steady state length distribution found in normally anchored rings, $f_{\mathrm{ss}}(l)$. (Note that the distribution evolves as the partially unanchored ring constricts, because cofilin-mediated severing is no longer balanced by actin polymerization in the permeabilized protoplast.) To compute $f_{\mathrm{ss}}(l)$, we denote the number of filaments in the ring with length between $l$ and $l+\Delta l$ as $F(l,t)\Delta l$. The dynamics of the length distribution are

$$\frac{\partial F(l,t)}{\partial t} = r_{\mathrm{nucl}}\delta(l) - v_{\mathrm{pol}}\frac{\partial F(l,t)}{\partial l} - r'_{\mathrm{sev}} l F(l,t) + r'_{\mathrm{sev}}\int_l^\infty F(l',t)\mathrm{d}l' - k'^{\mathrm{for}}_{\mathrm{off}}F(l,t) \quad (9)$$

where $k'^{\mathrm{for}}_{\mathrm{off}} = 0.023\ \mathrm{s}^{-1}$, $r'_{\mathrm{sev}} = 1.8\ \mu\mathrm{m}^{-1}\mathrm{min}^{-1}$ and $v_{\mathrm{pol}} = 70$ nm/s are the formin off rate, actin severing rate per filament length by cofilin and formin-mediated barbed end actin polymerization rate in an intact cell, respectively (3). The first term on the right hand side represents nucleation (with nucleation rate $r_{\mathrm{nucl}}$), the second term polymerization of actin subunits, the third and fourth terms cofilin severing, and the fifth term unbinding of formin from the ring.

Setting $\partial F(l,t)/\partial t = 0$ at steady state, and taking $\partial/\partial l$ of both sides, we have, for $l > 0$:

$$0 = -v_{\mathrm{pol}}\frac{\partial^2 F(l,t)}{\partial l^2} - \left(r'_{\mathrm{sev}} l + k'^{\mathrm{for}}_{\mathrm{off}}\right)\frac{\partial F(l,t)}{\partial l} - 2r'_{\mathrm{sev}}F(l,t) \quad (10)$$

Solving this equation and normalizing the total probability to unity yields the steady state actin filament length distribution $f_{\mathrm{ss}}(l)$:

$$f_{\mathrm{ss}}(l) = \left[\frac{k'^{\mathrm{for}}_{\mathrm{off}} + l\, r'_{\mathrm{sev}}}{v_{\mathrm{pol}}}\right]\exp\left[-\frac{l\left(2k'^{\mathrm{for}}_{\mathrm{off}} + l\, r'_{\mathrm{sev}}\right)}{2v_{\mathrm{pol}}}\right] \quad (11)$$

with $\int_0^\infty f_{\mathrm{ss}}(l)\,\mathrm{d}l = 1$, and the mean actin filament length $\langle l\rangle_{f_{\mathrm{ss}}} = \int_0^\infty l f_{\mathrm{ss}}(l)\,\mathrm{d}l = 1.3\ \mu$m.

In the experiments of Mishra et al., upon addition of ATP one (or possibly more) regions of the ring pulled away from the membrane within ~ 10 s, but neither the detailed dynamics of this detaching episode nor the shape of the unanchored segment immediately following detachment were reported (2). Therefore we did not attempt to describe the detachment episode itself, but instead we begin simulations at the instant immediately following detachment. To account for the detachment process, simulations used an initial condition in which a section of the ring is anchored and lies along the inner surface of the plasma membrane, a sphere of radius $R$, while the remainder of the ring (the unanchored segment) is slightly displaced in the inward radial direction from the spherical membrane surface (Fig. 2b), as follows. We use cylindrical coordinates $(r,\phi,z)$ with the origin at the center of the cell ghost, such that a ring initially lies in the x-y plane. We identify ring components in the unanchored segment by their azimuthal coordinates $\phi$ in the range ($0 < \phi < l_{\mathrm{u}}/R$), where $l_{\mathrm{u}}$ is the length of the unanchored segment and $R$ is the radius of the ring. Their radial locations $r$ are slightly decreased by a factor $f(\phi) < 1$, such that the post-detachment coordinates are $(r \times f(\phi), \phi, z)$. This factor $f(\phi)$ is a smooth random function of the azimuthal coordinate $\phi$ that has values between 0.9 and 1 for $0 < \phi < l_{\mathrm{u}}/R$, and is equal to 1 at $\phi = 0$ and $\phi = l_{\mathrm{u}}/R$.

Due to the random nature of the ring component distributions and the random nature of the displacement of the unanchored segment in the initial condition, rings occasionally snapped or became highly twisted. These simulations were discarded, since such phenomena were not reported in ref. (2).

#### b. Computation Scheme

Given an existing configuration, i.e. the positions of all the molecules in the simulation, we write force balance equations with both velocities and forces as unknown variables. This scheme is adapted from the method outlined in Witkin et al. (23). The force balance equations are:

$$
\begin{cases}
\sum_\alpha \left[\boldsymbol{f}_{i,\alpha}^{\mathbf{cap}} + \boldsymbol{f}_{i,\alpha}^{\mathbf{pull}}(\boldsymbol{v}_i, \boldsymbol{v}_\alpha)\right] - \frac{\partial H^{\mathrm{B}}}{\partial \boldsymbol{r}_i} + f_{i,i+1}^{\mathrm{tension}} \frac{\boldsymbol{r}_{i+1}-\boldsymbol{r}_i}{|\boldsymbol{r}_{i+1}-\boldsymbol{r}_i|} + \boldsymbol{f}_i^{\mathbf{mb}} + f_i^{\mathrm{anch}}\hat{r} + \boldsymbol{f}_{\mathbf{for}}^{\mathbf{drag,mb}}(\boldsymbol{v}_i) = 0, \\
\text{for anchored formin } i \\
\sum_\alpha \left[\boldsymbol{f}_{i,\alpha}^{\mathbf{cap}} + \boldsymbol{f}_{i,\alpha}^{\mathbf{pull}}(\boldsymbol{v}_i, \boldsymbol{v}_\alpha)\right] - \frac{\partial H^{\mathrm{B}}}{\partial \boldsymbol{r}_i} + f_{i,i+1}^{\mathrm{tension}} \frac{\boldsymbol{r}_{i+1}-\boldsymbol{r}_i}{|\boldsymbol{r}_{i+1}-\boldsymbol{r}_i|} + \boldsymbol{f}_i^{\mathbf{mb}} + \boldsymbol{f}_{\mathbf{for}}^{\mathbf{drag,bulk}}(\boldsymbol{v}_i) = 0, \\
\text{for unanchored formin } i \\
\sum_\alpha \left[\boldsymbol{f}_{i,\alpha}^{\mathbf{cap}} + \boldsymbol{f}_{i,\alpha}^{\mathbf{pull}}(\boldsymbol{v}_i, \boldsymbol{v}_\alpha)\right] - \frac{\partial H^{\mathrm{B}}}{\partial \boldsymbol{r}_i} + f_{i,i+1}^{\mathrm{tension}} \frac{\boldsymbol{r}_{i+1}-\boldsymbol{r}_i}{|\boldsymbol{r}_{i+1}-\boldsymbol{r}_i|} + f_{i-1,i}^{\mathrm{tension}} \frac{\boldsymbol{r}_{i-1}-\boldsymbol{r}_i}{|\boldsymbol{r}_{i-1}-\boldsymbol{r}_i|} + \boldsymbol{f}_i^{\mathbf{x}} + \boldsymbol{f}_i^{\mathbf{mb}} + \boldsymbol{f}_i^{\mathbf{visc}}(\boldsymbol{v}_i) = 0, \\
\text{for actin subunit } i \text{ not at the pointed end} \\
\sum_\alpha \left[\boldsymbol{f}_{i,\alpha}^{\mathbf{cap}} + \boldsymbol{f}_{i,\alpha}^{\mathbf{pull}}(\boldsymbol{v}_i, \boldsymbol{v}_\alpha)\right] - \frac{\partial H^{\mathrm{B}}}{\partial \boldsymbol{r}_i} + f_{i-1,i}^{\mathrm{tension}} \frac{\boldsymbol{r}_{i-1}-\boldsymbol{r}_i}{|\boldsymbol{r}_{i-1}-\boldsymbol{r}_i|} + \boldsymbol{f}_i^{\mathbf{x}} + \boldsymbol{f}_i^{\mathbf{mb}} + \boldsymbol{f}_i^{\mathbf{visc}}(\boldsymbol{v}_i) = 0, \\
\text{for actin subunit } i \text{ at the pointed end} \\
\sum_i \left[-\boldsymbol{f}_{i,\alpha}^{\mathbf{cap}} - \boldsymbol{f}_{i,\alpha}^{\mathbf{pull}}(\boldsymbol{v}_i, \boldsymbol{v}_\alpha)\right] + \boldsymbol{f}_\alpha^{\mathbf{excl}} + \boldsymbol{f}_\alpha^{\mathbf{mb}} + f_\alpha^{\mathrm{anch}}\hat{r} + \boldsymbol{f}_{\mathbf{myo}}^{\mathbf{drag,mb}}(\boldsymbol{v}_\alpha) = 0, \text{for anchored myosin cluster } \alpha \\
\sum_i \left[-\boldsymbol{f}_{i,\alpha}^{\mathbf{cap}} - \boldsymbol{f}_{i,\alpha}^{\mathbf{pull}}(\boldsymbol{v}_i, \boldsymbol{v}_\alpha)\right] + \boldsymbol{f}_\alpha^{\mathbf{excl}} + \boldsymbol{f}_\alpha^{\mathbf{mb}} + \boldsymbol{f}_{\mathbf{myo}}^{\mathbf{drag,,bulk}}(\boldsymbol{v}_\alpha) = 0, \text{for unanchored myosin cluster } \alpha
\end{cases}
\quad (8)
$$

We then numerically solve for the unknown variables $\{\boldsymbol{v}_i\}$, $\{\boldsymbol{v}_\alpha\}$, $\{f_{i,i+1}^{\mathrm{tension}}\}$, $\{f_i^{\mathrm{anch}}\}$ and $\{f_\alpha^{\mathrm{anch}}\}$ from these equations. Given the velocities of ring components, the system is evolved using the Euler method with a time step $\Delta t = 1$ ms.

We found that when the time step exceeds a certain value, artificial oscillations occurred in the simulations of rapidly diverging amplitude. To allow for the use of a larger time step, we suppressed such oscillations using a pairwise drag $\boldsymbol{f} = -\gamma_{\mathrm{a}}\boldsymbol{\Delta v}$ between pairs of adjacent actin subunits and between interacting myosin cluster/actin subunit pairs, where $\boldsymbol{\Delta v}$ is the relative velocity and the artificial drag coefficient $\gamma_{\mathrm{a}} = 0.2\ \mathrm{pN \cdot s/\mu m}$ was chosen to allow for rapid computation. Being a pairwise force, this drag produces zero net force and does not affect the dynamics of the ring as a whole.

#### c. Calculation of ring length

The initial condition for the simulation includes the ring shape, and we lay down ring components along the ring contour that follows this shape. The ring components are then free to evolve, and the ring shape and length evolve with time as an *output* of the simulation. Mishra et al. measured ring length from fluorescence images of myosin-II light chain Rlc1p (2). Correspondingly, in the simulation we use the positions of all myosin-II clusters $\{\boldsymbol{r}_\alpha\}$ to generate a smooth ring contour. We perform a smoothing spline fit of the radial coordinates $\{\rho_\alpha\}$ and heights $\{z_\alpha\}$ of myosin-II clusters as a function of their azimuthal coordinates $\{\phi_\alpha\}$ (here the origin of the cylindrical coordinates is set at the mean myosin position $\langle \boldsymbol{r}_\alpha \rangle$). At selected time intervals during the simulation we select 200 points on the smooth contour of the ring evenly spaced in $\phi$, and sum the distance between adjacent points to obtain the length of the ring. In this way the length of the ring is evolved throughout the simulation.

#### d. Calculation of ring constriction rate

Given the ring length $L(t)$ as a function of time $t$ at discrete time points evenly spaced by $\Delta t = 1\ s$, we calculate the constriction rate $v(t) = -dL/dt$ with a fourth-order finite difference scheme:

$$v(t) = -[-L(t + 2\ \mathrm{s}) + 8L(t + 1\ \mathrm{s}) - 8L(t - 1\ \mathrm{s}) + L(t - 2\ \mathrm{s})]/12\ s \quad (12)$$

For the initial constriction rate, $v_0$, the above scheme cannot be applied. We use instead a second-order forward difference scheme:

$$v_0 = [-3L(1\ \mathrm{s}) + 4L(2\ \mathrm{s}) - L(3\ \mathrm{s})]/2\ \mathrm{s} \quad (13)$$

For time-averaged constriction rates, note that taking the arithmetic mean of constriction rates at all time points would give an incorrect result. Therefore we performed a least-squares fit of $L(t)$ as a function of time $t$, and took time-averaged constriction rates as the slope of the best-fit line.

##### e. Calculation of ring tension

For a bundle, tension is the attractive force between two segments of the bundle separated by an imaginary plane. For a bundle that interacts with its environment, e.g. the cytokinetic ring which interacts with other parts of the cell such as the plasma membrane, the tension is a local quantity whose value varies as one moves along the bundle. Indeed, in a partially unanchored ring as realized in the experiments of Mishra et al. (2), the anchored and unanchored segments have completely different interactions with the environment and the tension is essentially zero in the unanchored segment, but much higher $\sim 300$ pN in the anchored segment. We use the following method to numerically measure the local ring tension.

To calculate the local ring tension $T$ as a function of position $s$ along the ring, we calculate both tension $T(\phi)$ and position $s(\phi)$ as functions of the azimuthal coordinate $\phi$ (here the origin of the cylindrical coordinates is set at the mean myosin position $\langle \boldsymbol{r}_\alpha \rangle$). For the tension $T(\phi)$, we choose 200 imaginary planes $\phi_n = n \cdot (2\pi/200)$ evenly spaced in $\phi$. $T(\phi_n)$ is calculated as the sum of pairwise attractive forces $\boldsymbol{f}_{i,i+1}^{\mathbf{tension}}$ between all actin subunit pairs whose connecting rod is intersected by the plane $\phi_n$, projected perpendicular onto the plane $\phi_n$. For the positions $s(\phi)$, we selected $\phi_1$ as the starting point and calculate $s(\phi_n)$ as the distance along the ring up to $\phi_n$, by cumulatively summing the distance between adjacent points $(\phi_1, \phi_2)$, $(\phi_2, \phi_3)$, …, $(\phi_{n-1}, \phi_n)$ on the ring contour (Sec. 2c). We then use $T(\phi_n)$ and $s(\phi_n)$ to obtain the relation $T(s)$.

##### f. Calculation of ring component densities

In Fig. 4c of the main text we present ring component densities, namely, the number of actin filaments in the cross section and the myosin density (number of heads per unit length). The densities are shown as functions of position $s$ along the ring at time t = 5 s, averaged over n = 7 rings. The averaging procedure is as follows. We use cylindrical coordinates $(r, \phi, z)$ with the origin at the center of the cell ghost, such that a ring initially lies in the x-y plane. We chose a large number of evenly spaced $\phi$ values, and we calculate the ring component densities and $s(\phi)$ at each of these locations, where $s(\phi)$ is the distance along the ring at $\phi$, measured from $\phi = 0$. This is repeated for 7 rings and the average density at each $s$ value is plotted.

#### 3. Determination of model parameters

In this section we describe how the values of several model parameters were determined. For other parameters, see Table S1.

##### a. Determination of myosin-II load-free velocity $v_{\mathbf{myo}}^{\mathbf{0}}$ from the motility assay of ref. (1)

In their *in vitro* motility assays, Stark et al. (1) measured the gliding velocity of tropomyosin Cdc8p-associated actin filaments as a function of the number of Myo2p heads interacting with each filament. All proteins were purified from fission yeast. Now previous experiments have shown that in normal intact yeast cells there are ~ 3000 Myo2p and ~ 150 formin Cdc12 dimers in the ring (7), and our model assumes every formin dimer in the ring caps an actin filament, and every actin filament in the ring is capped by a formin dimer. Thus in normal cells there are ~ 150 actin filaments in the ring, so ~ 20 Myo2p heads interact with each actin filament. We assume that the number of Myo2p and Cdc12p per unit length along the ring is the same in intact cells and in protoplasts, so again ~ 20 Myo2p heads interact with each actin filament in protoplasts. Using this value, the measurements by Stark et al. (1) yield a value $v_{\rm myo}^{0} = 0.24\ \mu{\rm m/s}$ for the myosin-II load-free velocity.

Note this argument assumes that all Myo2p heads interact with an actin filament (there are no idle heads), i.e. myosin-II clusters in the ring are saturated with actin filaments. We argue that this is indeed the case, as there are more available actin binding sites among all the filaments than there are myosin-II heads (3). These saturation effects are an important feature of the model (see "*Myosin-II saturation effects*" above).

#### b. Turnover parameters

Turnover kinetics of a number of ring components have been measured in fission yeast (3, 9, 10). However, these kinetics are dramatically different in permeabilized protoplasts.

##### (i) Myosin-II cluster off rate, $k_{\rm off}^{\rm myo}$

In the experiments of Mishra et al. (2), 78% of the myosin-II regulatory light chain Rlc1p initially in the permeablized protoplast ring was still present once constriction was complete, as measured by Western blot analysis. Taking a constriction time of 60 s (2), we obtain the myosin-II cluster off rate $k_{\rm off}^{\rm myo} = -\log(0.78)/(60\ {\rm s}) = 0.0041\ {\rm s}^{-1}$. Interestingly, this value is ~ 6-fold smaller than the value in normal cells (3, 9).

##### (ii) Formin off rate, $k_{\rm off}^{\rm for}$

Mishra et al. (2) found that when the F-actin-stabilizing drug jasplakinolide was added, 73% of the actin initially present in the ring was still present after constriction, as measured by Western blot analysis. We assumed that jasplakinolide completely protected the actin filaments so that all actin lengths were constant in time, and actin could only exit the ring by dissociation of whole filaments when formins dissociated (Fig. 2a). Thus the fraction of actin remaining in the ring equals the fraction of formin remaining in the ring. Taking the duration of constriction as 60 s (2), we obtain the formin off rate $k_{\rm off}^{\rm for} = -\log(0.73)/(60\ {\rm s}) = 0.0052\ {\rm s}^{-1}$. Interestingly, this value is ~ 4-fold smaller than the value in normal cells (3, 10).

##### (iii) Actin severing rate per filament length by cofilin, $r_{\rm sev}$

Now that we have obtained the formin off rate $k_{\rm off}^{\rm for}$, we use it to estimate the remaining actin turnover parameter, the actin severing rate per filament length by cofilin, $r_{\rm sev}$. Given the initial distribution of filament lengths $f(l, t = 0) = f_{\rm ss}(l)$ in the ring, which we assume to follow the steady state length distribution $f_{\rm ss}(l)$ in normally anchored rings (see Sec. 2a), we compute distribution $f(l,t)$ and the decrease in mean filament length $\langle l(t)\rangle_f / \langle l \rangle_{f_{\rm ss}}$ at a later time $t$. Multiplying this with the fraction of formin dimers that remain in the ring after constriction, we obtain the fraction of actin that remains in the ring at time $t$. We compare this to Mishra et al.'s observation that (in the absence of jasplakinolide) 47% of actin remains in the ring after constriction (2), to obtain $r_{\rm sev}$. Detailed procedure follows.

*Mean length versus time of filaments with uniform initial length $l_0$.* Given a large number of actin filaments with the same initial length $l_0$ at $t = 0$, at a later time $t$ they will have different lengths $l_i(t)$ due to the randomness of cofilin severing. Let the length distribution be $g(l,t)$, where $\int_0^{l_0} g(l,t)dl = 1$ and $g(l,0) = \delta(l - l_0)$. We stress that $g(l,t)$ is not to be confused with $f(l,t)$, which begins with initial length distribution $f_{\mathrm{ss}}(l_0)$.

If no severing has occurred between the barbed end $x = 0$ and some point $x_1 < l_0$ on the $i$th filament, the filament length $l_i(t)$ is at least $x_1$. Mathematically, this means

$$\exp(-r_{\mathrm{sev}}\, x_1 t) = \int_{x_1}^{l_0} g(l,t)dl \quad (14)$$

where the LHS is the probability that no severing events occur between $x = 0$ and $x = x_1$ up to time $t$ on a given filament, and the RHS is the probability that this filament has length at least $x_1$. Taking the derivative with respect to $x_1$ on both sides, we have $g(x_1,t) = r_{\mathrm{sev}}\, t \exp(-r_{\mathrm{sev}}\, x_1\, t)$ for all $x_1 < l_0$. Together with the probability $e^{-r_{\mathrm{sev}}\, l_0\, t}$ that no severing events occur on the whole filament (leaving its length intact at $l_i(t) = l_0$, we have

$$g(l,t) = r_{\mathrm{sev}}\, t\, e^{-r_{\mathrm{sev}}\, l\, t} + \delta(l - l_0)e^{-r_{\mathrm{sev}}\, l_0\, t} \quad (15)$$

$$\langle l(t)\rangle_g = \int_0^{l_0} l\, g(l,t)dl = l_0 \frac{1 - e^{-r_{\mathrm{sev}}\, l_0\, t}}{r_{\mathrm{sev}} l_0 t} \quad (16)$$

Where the variable $x_1$ has been replaced by $l$, and the subscript $g$ denotes the ensemble of filaments starting with a uniform initial length $l_0$.

*Mean length versus time of filaments with initial length distribution $f_{ss}(l_0)$.* Given a large number of actin filaments with initial length distribution $f(l_0, t = 0) = f_{ss}(l_0)$, we can group the filaments according to their initial length $l_0$, and the mean length versus time of each group with initial length $l_0$ is given by Eq. 16. Therefore, the mean length at a later time $t$ for the entire ensemble is

$$\langle l(t)\rangle_f = \int_0^{\infty} l_0 \frac{1 - e^{-r_{\mathrm{sev}}\, l_0\, t}}{r_{\mathrm{sev}} l_0 t} f_{\mathrm{ss}}(l_0) dl_0 \quad (17)$$

At the end of constriction, which we estimate as $t = 60$ s, the fraction of actin remaining in the ring is $\exp(-k_{\mathrm{off}}^{\mathrm{for}}\, t) \cdot \langle l(t)\rangle_f / \langle l\rangle_{f_{\mathrm{ss}}}$. In experiment, Mishra et al. reported that in the absence of jasplakinolide 45% of actin remains in the ring after constriction (2). Therefore, we numerically solved the equation $\exp(-k_{\mathrm{off}}^{\mathrm{for}}\, t) \cdot \langle l(t)\rangle_f / \langle l\rangle_{f_{\mathrm{ss}}} = 0.45$ and obtained $r_{\mathrm{sev}} = 0.011\ \mu\mathrm{m}^{-1}\mathrm{s}^{-1}$.

#### 4. The reeling-in constriction mechanism

Simulations of our model revealed that unanchored segments shortened by being reeled in at their two ends where they join the anchored segment. Here we discuss the reeling-in mechanism in some detail.

The agents of reeling in are *anchored actin filaments in the interfacial zone* (i.e., the location where the anchored and unanchored segments meet) that straddle the interface. These filaments have barbed ends anchored on the anchored segment side of the interface, and their pointed ends extend into the unanchored segment (Fig. 5c). Note that, among filaments that are anchored, any filament that straddles this interface is bound to have this particular polarity. While other filaments straddle the interface with the opposite polarity (pointed ends extending into the anchored segment) all such filaments are unanchored. In other words, there is a 100% polarity bias of anchored actin filaments at the interface. This is entirely a consequence of the barbed end anchoring scheme that is assumed. As one moves into the anchored segment away from the interface this bias weakens (there is no polarity bias at the center of the anchored segment.)

The motion of unanchored myosin-II clusters in the unanchored ring segment is dominated by anchored actin filaments that they interact with. Near the interface, anchored filaments of the same orientation therefore reel the unanchored myosin-II clusters towards their barbed ends, that is, into the anchored segment (Fig. 5c). (There is a very small influence from unanchored actin filaments which slide, relative to myosin, at close to the load-free velocity, $v_{\mathrm{myo}}^{0}$, against almost zero drag force.)

If one ignores both the sliding resistance from anchored myosin clusters on incoming actin filaments, and myosin crowding effects at the interface (to be discussed later), the speed that unanchored myosin-II clusters move on anchored actin filaments is the load-free velocity $v_{\mathrm{myo}}^{0}$, giving a shortening rate $2v_{\mathrm{myo}}^{0}$ of the unanchored segment due to the contributions at both interfaces (Fig. 5a). The same result can be obtained if one calculates the shortening rate of the unanchored segment based on the motion of actin filaments, as follows. Consider all unanchored filaments with one orientation: at one interface, they move into the anchored segment at $2v_{\mathrm{myo}}^{0}$, being propelled at $v_{\mathrm{myo}}^{0}$ relative to unanchored myosin-II clusters which are themselves moving at $v_{\mathrm{myo}}^{0}$ (Fig. 5c); however at the other interface they have almost zero velocity, being propelled away from the anchored segment at velocity $v_{\mathrm{myo}}^{0}$ relative to unanchored myosin-II clusters which are moving at $v_{\mathrm{myo}}^{0}$ into the anchored segment. Therefore, the shortening rate of the unanchored segment is $2\ v_{\mathrm{myo}}^{0}$ no matter it is calculated based on myosin cluster motion or actin filament motion.

Relative to the expectation of the above simplified argument, two sources of resistance slow down the reeling-in velocity to a value $\sim 0.5v_{\mathrm{myo}}^{0}$ (Fig. 4b), giving a constriction rate $\sim v_{\mathrm{myo}}^{0}$ (not $\sim 2\ v_{\mathrm{myo}}^{0}$). First, if an actin filament entered the anchored segment with velocity $2v_{\mathrm{myo}}^{0}$ it would have velocity $\sim 2v_{\mathrm{myo}}^{0}$ relative to the slow-moving anchored myosin-II clusters. Due to the linear myosin force-velocity relation such a velocity, being greater than $v_{\mathrm{myo}}^{0}$, would result in a reverse myosin force resisting the motion of the actin filament. Therefore the actual velocity of an actin filament being reeled in lies between $v_{\mathrm{myo}}^{0}$ and $2v_{\mathrm{myo}}^{0}$ (Fig. 4e), reflecting a tug-of-war between the reeling-in mechanism that tends to reel in filaments at $2v_{\mathrm{myo}}^{0}$ and the anchored myosin-II clusters that tend to slide the filaments in at $v_{\mathrm{myo}}^{0}$ (and resist the motion of filaments with velocities that exceed $v_{\mathrm{myo}}^{0}$).

Second, due to the non-contractile reeling in, myosin accumulates in puncta of growing amplitude near the anchored/unanchored interfaces (Fig. 4a,c, main text). The crowding of myosin-II clusters at the interface, due to their finite size and high density, tends to block other incoming myosin-II clusters. This blockage is not static, however, as it consists of unanchored myosin-II clusters and is thus constantly pushed into the anchored segment by incoming myosin-II clusters. The incoming myosin-II clusters that would have moved at $v_{\mathrm{myo}}^{0}$ are slowed down by this crowding of myosin at the interface.

### Supplementary Note 2

### Fitting the model-predicted ATP-dependence of the myosin-II load-free velocity to Michaelis-Menten kinetics

We fit the calculated the ATP-dependence of $v_{\mathrm{myo}}^{0}$ (Fig. 7 of main text) to Michaelis-Menten kinetics

$$v_{\mathrm{myo}}^{0} = \frac{v_{\mathrm{max}}\,[\mathrm{ATP}]}{K_{\mathrm{M}} + [\mathrm{ATP}]}$$

with a nonlinear least-squares method. Here $v_{\mathrm{max}}$ is the load-free velocity at saturating ATP concentration, [ATP] is the ATP concentration and $K_{\mathrm{M}}$ is the Michaelis constant, namely the ATP concentration at which a half-maximal load-free velocity is reached. Our fitting procedure yielded $v_{\mathrm{max}} = 0.23$ µm $\mathrm{s}^{-1}$, close to the 0.24 µm/s reported in ref. (1), and $K_{\mathrm{M}} = 30$ µM.

Note that, due to the linear dependence of the ring constriction rate on the myosin-II load-free velocity that has emerged from this study (Fig. 3j), the Michaelis constant for the load-free velocity turns out to be almost equal to the analogous quantity reported by Mishra et al. (2) for the constriction rate itself, namely $\widehat{K}_M = 32$ µM. However, the significance of these two values is dramatically different: $K_{\mathrm{M}}$ for the load-free velocity characterizes the enzyme kinetics of fission yeast myosin-II, while $\widehat{K}_M$ for the ring constriction rate merely parametrizes a phenomenological relation. In fact, our study has shown that the origin of the constriction rate appearing to follow Michaelis-Menten kinetics is the constriction rate's very particular *linear* dependence on the myosin-II load-free velocity.

### Movies

**Movie S1**

Simulated constriction of a partially unanchored ring. The unanchored segment shortens, while the anchored segment remains almost constant in length. Simulation parameters as for Fig. 3. Red: actin filaments. Cyan: plasma membrane. Thicknesses of actin filaments and of the plasma membrane are not to scale.

**Movie S2**

Simulated component motions in the anchored segment of a partially unanchored ring, projected onto the plane of ring constriction. Simulation parameters as for Fig. 3. Red: actin filaments. Blue: formin dimers. Orange: myosin-II clusters. The plasma membrane is implemented as a potential limiting the range of ring component motions (see Methods). Ring components are not depicted to scale.

**Movie S3**

The unanchored ring segment shortens by being reeled in by the anchored segment. Simulated component motions are shown in the vicinity of the anchored-unanchored interface (dashed line) for a partially unanchored ring, projected onto the plane of ring constriction. Simulation parameters as for Fig. 3. There is a steady flow of ring components into the anchored segment: unanchored myosin-II clusters (orange), unanchored actin filaments (red, not resolved in this video) and formin dimers (cyan) belonging to the barbed ends of unanchored actin filaments. Anchored myosin-II clusters (brown) and their formin dimers (blue) move slowly. The interface is defined as the location of the last anchored myosin or formin at the edge. Ring components are not depicted to scale.

**Movie S4**

Completely unanchored rings do not constrict. Simulated evolution of a completely unanchored ring. Simulation parameters as for Fig. 3. Red: actin filaments. Thicknesses of actin filaments and the plasma membrane are not to scale.

**Movie S5**

Partially anchored rings fail to constrict when the actin filaments in the anchored segment are not anchored. Simulated evolution of a partially anchored ring, with unanchored actin filaments and anchored myosin-II clusters in the anchored segment. Simulation parameters as for Fig. 3. Red: actin filaments. Thicknesses of actin filaments and the plasma membrane are not to scale.

**Movie S6**

Partially anchored rings constrict even when myosin-II clusters in the anchored segment are unanchored. Simulated evolution of a partially anchored ring, with unanchored myosin-II clusters and anchored actin filaments in the anchored segment. Simulation parameters as for Fig. 3. Red: actin filaments. Thicknesses of actin filaments and the plasma membrane are not to scale.

### Table S1. Model parameters

| Parameter | Meaning | Value | Legend |
|---|---|---|---|
| $\rho_{\mathrm{for}}$ | Initial mean density of formin dimers along the ring | $15\ \mu\mathrm{m}^{-1}$ | (A) |
| $\rho_{\mathrm{myo}}$ | Initial mean density of myosin-II clusters along the ring | $18.75\ \mu\mathrm{m}^{-1}$ | (B) |
| $r_{\mathrm{sev}}$ | Cofilin severing rate of actin filaments | $0.011\ \mu\mathrm{m}^{-1}\mathrm{s}^{-1}$ | (C) |
| $k_{\mathrm{off}}^{\mathrm{for}}$ | Formin off rate | $0.0052\ \mathrm{s}^{-1}$ | (C) |
| $k_{\mathrm{off}}^{\mathrm{myo}}$ | Myosin-II cluster off rate | $0.0041\ \mathrm{s}^{-1}$ | (C) |
| $k_{\mathrm{off}}^{\mathrm{x}}$ | α-actinin off rate | $3.3\ \mathrm{s}^{-1}$ | (D) |
| $f_{\mathrm{s}}$ | Myosin-II cluster stall force | 4 pN | (E) |
| $v_{\mathrm{myo}}^{0}$ | Myosin-II load-free velocity | $0.24\ \mu\mathrm{m/s}$ | (F) |
| $r_{\mathrm{myo}}$ | Myosin-II cluster capture radius for actin filaments | 0.1 μm | (G) |
| $l_{\mathrm{p}}$ | Actin filament persistence length | 10 μm | (H) |
| $\gamma_{\mathrm{myo}}$ | Myosin-II cluster anchor drag coefficient | $0.52\ \mathrm{nN}\cdot\mathrm{s}/\mu\mathrm{m}$ | (I) |
| $\gamma_{\mathrm{for}}$ | Formin anchor drag coefficient | $1.9\ \mathrm{nN}\cdot\mathrm{s}/\mu\mathrm{m}$ | (E) |

(A) Ref. (7).

(B) Obtained by dividing the density of Myo2p myosin-II heavy chains (~3000 Myo2p myosin-II heavy chains in a ring of ~10 μm in length, Ref. (7)) by 16 heavy chains per cluster.

(C) Determined from the experimental results of Ref. (2). See Supplementary Note 1.

(D) Refs. (16), (17), (18).

(E) From the measurement of node motions in Ref. (3).

(F) Estimated from Ref. (1). See Methods.

(G) Estimated from single-molecule high resolution colocalization (SHREC) measurements of the distance that myosin heads extend from precursor nodes (Ref. (8)).

(H) Refs. (13), (24).

(I) Ref. (3) reported $1.3\ \mathrm{nN}\cdot\mathrm{s}/\mu\mathrm{m}$. In the present study, 40% of this value is used because the myosin-II cluster size is 40% of the size assumed in Ref. (3) (16 heads versus 40 heads).

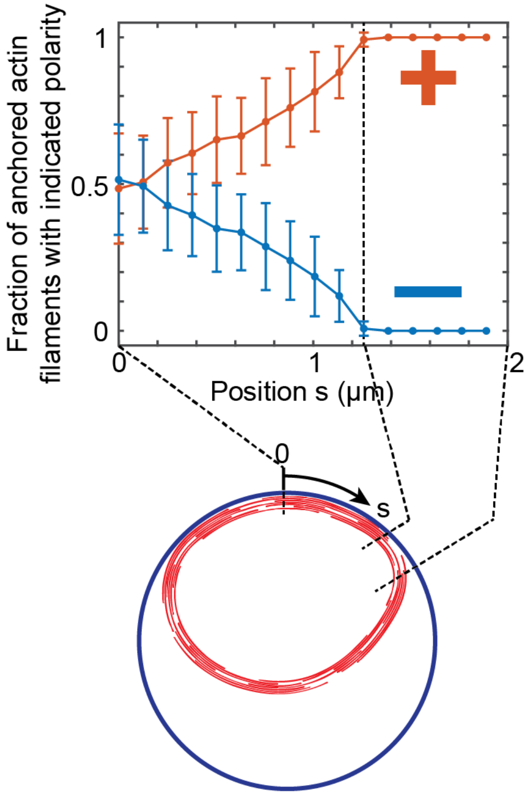


**Fig. S1.** Anchored filaments that straddle the anchored/unanchored interface all have the same polarity, with their pointed ends orienting into the unanchored region. At each location, the polarities of all *anchored* filaments passing through that location were recorded in simulations with parameters as for Fig. 3. The fraction with each polarity is plotted versus distance, averaged over n = 10 rings at time 10 s ("+" and "-" denote clockwise and anticlockwise polarities, respectively). Near the interface, almost all anchored filaments have clockwise polarity. Error bars: s.d.

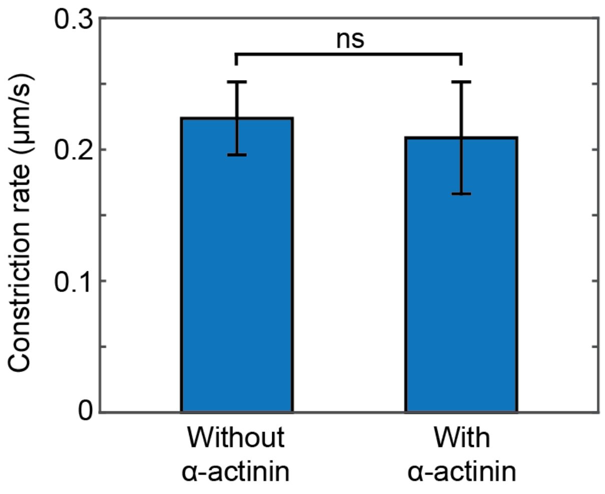


**Fig. S2.** α-actinin crosslinking has no effect on the ring constriction rate. Model parameters as for Fig. 3 of main text. There is no statistically significant difference between the time-averaged constriction rate of simulated rings without α-actinin ($n = 10$) and with α-actinin ($n = 7$), $p = 0.40$.

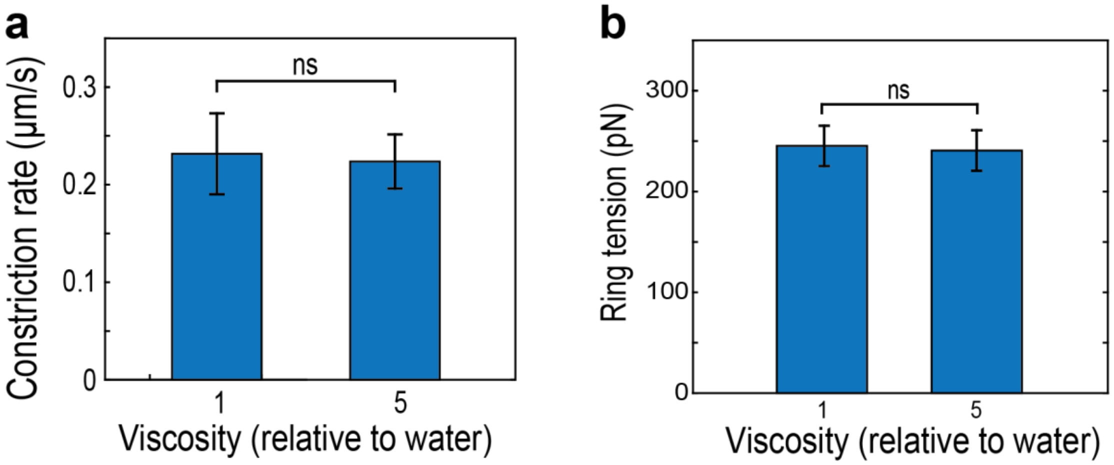


**Fig. S3.** The simulated ring tension and constriction rate are unaffected by the value of the viscosity of the aqueous medium used in the simulation. Other model parameters, as in Fig. 3 of main text. (**a**) Time-averaged constriction rate using a value of the viscosity equal to that of water (n = 8) and 5 times that of water (n = 10). (**b**) Mean tension of the anchored segment of the cytokinetic ring in simulations using a value of the viscosity equal to that of water (n = 8) and 5 times that of water (n = 10). Error bars: s.d.

### References


1. Stark BC, Sladewski TE, Pollard LW, & Lord M (2010) Tropomyosin and myosin-II cellular levels promote actomyosin ring assembly in fission yeast. *Mol. Biol. Cell* 21(6):989-1000.
2. Mishra M*, et al.* (2013) In vitro contraction of cytokinetic ring depends on myosin II but not on actin dynamics. *Nat. Cell Biol.* 15(7):853-859.
3. Stachowiak MR*, et al.* (2014) Mechanism of cytokinetic contractile ring constriction in fission yeast. *Dev. Cell* 29(5):547-561.
4. Matsuyama A*, et al.* (2006) ORFeome cloning and global analysis of protein localization in the fission yeast Schizosaccharomyces pombe. *Nat. Biotechnol.* 24(7):841-847.
5. Arasada R & Pollard TD (2014) Contractile ring stability in S. pombe depends on F-BAR Protein Cdc15p and Bgs1p transport from the Golgi complex. *Cell Rep.* 8(5):1533-1544.
6. Goss JW, Kim S, Bledsoe H, & Pollard TD (2014) Characterization of the roles of Blt1p in fission yeast cytokinesis. *Mol. Biol. Cell* 25(13):1946-1957.
7. Wu JQ & Pollard TD (2005) Counting cytokinesis proteins globally and locally in fission yeast. *Science* 310(5746):310-314.
8. Laporte D, Coffman VC, Lee IJ, & Wu JQ (2011) Assembly and architecture of precursor nodes during fission yeast cytokinesis. *J. Cell Biol.* 192(6):1005-1021.
9. Pelham RJ & Chang F (2002) Actin dynamics in the contractile ring during cytokinesis in fission yeast. *Nature* 419(6902):82-86.
10. Yonetani A*, et al.* (2008) Regulation and targeting of the fission yeast formin cdc12p in cytokinesis. *Mol. Biol. Cell* 19(5):2208-2219.
11. Vavylonis D, Wu JQ, Hao S, O'Shaughnessy B, & Pollard TD (2008) Assembly mechanism of the contractile ring for cytokinesis by fission yeast. *Science* 319(5859):97-100.
12. Naqvi NI, Eng K, Gould KL, & Balasubramanian MK (1999) Evidence for F-actin-dependent and - independent mechanisms involved in assembly and stability of the medial actomyosin ring in fission yeast. *EMBO J.* 18(4):854-862.
13. Ott A, Magnasco M, Simon A, & Libchaber A (1993) Measurement of the persistence length of polymerized actin using fluorescence microscopy. *Phys. Rev. E* 48(3):R1642-R1645.
14. Kanbe T, Kobayashi I, & Tanaka K (1989) Dynamics of cytoplasmic organelles in the cell cycle of the fission yeast Schizosaccharomyces pombe: three-dimensional reconstruction from serial sections. *J. Cell. Sci.* 94:647-656.
15. Kojima H, Ishijima A, & Yanagida T (1994) Direct Measurement of Stiffness of Single Actin-Filaments with and without Tropomyosin by in-Vitro Nanomanipulation. *Proc. Natl. Acad. Sci. USA* 91(26):12962-12966.
16. Kuhlman PA, Ellis J, Critchley DR, & Bagshaw CR (1994) The Kinetics of the Interaction between the Actin-Binding Domain of Alpha-Actinin and F-Actin. *FEBS Lett.* 339(3):297-301.
17. Miyata H, Yasuda R, & Kinosita K (1996) Strength and lifetime of the bond between actin and skeletal muscle alpha-actinin studied with an optical trapping technique. *Biochim. Biophys. Acta.* 1290(1):83-88.
18. Xu JY, Wirtz D, & Pollard TD (1998) Dynamic cross-linking by alpha-actinin determines the mechanical properties of actin filament networks. *J. Biol. Chem.* 273(16):9570-9576.
19. Tyska MJ*, et al.* (1999) Two heads of myosin are better than one for generating force and motion. *Proc. NatL. Acad. Sci. USA* 96(8):4402-4407.
20. Gauger E & Stark H (2006) Numerical study of a microscopic artificial swimmer. *Phys. Rev. E* 74(2).

21. Fowler WE & Aebi U (1983) A Consistent Picture of the Actin Filament Related to the Orientation of the Actin Molecule. *J. Cell Biol.* 97(1):264-269.

22. Broersma S (1960) Viscous Force Constant for a Closed Cylinder. *J. Chem. Phys.* 32(6):1632.

23. Witkin A, Gleicher M, & Welch W (1990) Interactive dynamics. *Proceedings of the 1990 symposium on interactive 3D graphics*:11-21.

24. Riveline D, Wiggins CH, Goldstein RE, & Ott A (1997) Elastohydrodynamic study of actin filaments using fluorescence microscopy. *Phys. Rev. E* 56(2):R1330-R1333.
